# The cryo-EM structure of an adaptor-effector complex reveals the mechanism of a widespread pore-forming toxin family

**DOI:** 10.1101/2025.07.04.663240

**Authors:** Carmen Velázquez, Maialen Zabala-Zearreta, Carmen Paredes, Cristina Civantos, Jon Altuna-Alvarez, Patricia Bernal, David Albesa-Jové

**Author notes:** These authors contributed equally to this work (Carmen Velázquez, Maialen Zabala-Zearreta).

## Abstract

*Pseudomonas putida* KT2440 is a plant-beneficial rhizobacterium that encodes multiple Type VI secretion systems (T6SS) to outcompete phytopathogens in the rhizosphere. Among its antibacterial effectors, Tke5 has been identified as a potent pore-forming toxin that disrupts ion homeostasis without causing considerable membrane damage. Tke5 belongs to the BTH_I2691 protein family and harbours an N-terminal <u>m</u>arker for the type s<u>ix</u> secretion system effectors (MIX) motif, previously shown to be required for T6SS-dependent secretion in other systems. Many MIX-containing effectors require <u>T</u>6SS <u>a</u>daptor <u>p</u>roteins (Tap) for secretion, but until now, the molecular mechanism for adaptor-effector binding has remained elusive. Here, we report the 2.8 Å cryo-EM structure of the Tap3–Tke5 complex, providing structural and functional insight into how this effector is recruited by its cognate adaptor protein Tap3. Functional dissection shows that the α-helical region of Tke5 is sufficient to kill intoxicated bacteria, while its β-rich region likely contributes to target membrane specificity. These findings suggest a general mechanism of MIX-containing BTH_I2691 proteins for Tap recruitment and toxin activity, contributing to our fundamental understanding of a widespread yet understudied toxin family.

## Introduction

*Pseudomonas putida* KT2440 is a plant growth–promoting rhizobacterium of major interest in agricultural biotechnology due to its stimulatory and protective effects on plants. This strain exhibits several beneficial traits, including the production of indole compounds and siderophores, phosphate solubilisation, and 1-aminocyclopropane-1-carboxylate (ACC) deaminase activity. These features have been shown to enhance germination rates, increase root and shoot length, improve fresh and dry biomass, and trigger induced systemic resistance in plants (Costa-Gutierrez *et al*, 2022).

*P. putida* KT2440 harbours three distinct Type VI secretion system (T6SS) gene clusters, designated K1-, K2-, and K3-T6SS (Bernal *et al*, 2017). The T6SS is a sophisticated contractile nanomachine employed by many Gram-negative bacteria to deliver toxic effector proteins into neighbouring cells, thereby providing a competitive advantage in complex microbial communities (Mougous *et al*, 2006). In *P. putida* KT2440, the K1-T6SS has been shown to effectively eliminate a broad range of bacterial competitors, including recalcitrant phytopathogens, thereby playing a key role in preventing plant infections and enhancing the bacterium’s biocontrol activity (Bernal *et al*, 2017, 2021).

Among the effectors encoded within the K3-T6SS cluster of *P. putida* KT2440, a recent study identified a <u>T</u>ype six <u>K</u>T2440 <u>e</u>ffector 5 (Tke5) as a potent antibacterial toxin effective against ten of the most resilient plant pathogens (Velázquez *et al*, 2025). Notably, this work also elucidated Tke5’s molecular mechanism of action, showing that it functions as a novel pore-forming toxin (PFT) that kills target cells through selective ion transport. Instead of causing extensive membrane disruption, Tke5 induces membrane depolarization by perturbing ion homeostasis, ultimately leading to bacterial cell death while preserving overall membrane integrity (Velázquez *et al*, 2025). This work represented the first biophysical dissection for a member of the T6SS effector BTH_I2691 protein family (InterPro accession NF041559). Members of this family—including Tke5—harbour an N-terminal <u>m</u>arker for type s<u>ix</u> secretion system effectors (MIX) motif that was shown to be necessary for T6SS-dependent secretion (Salomon *et al*, 2014).

The genomic context of *tke5* within the K3-T6SS cluster is of particular interest. Near the 3’ end of this cluster, we find genes coding for a <u>V</u>aline-<u>g</u>lycine repeat protein <u>G</u> (VgrG3), a <u>T</u>ype VI <u>a</u>daptor <u>p</u>rotein (Tap3), Tke5, its cognate <u>T</u>ype six <u>K</u>T2440 immunity protein (Tki5), and a <u>T</u>ype <u>s</u>ix <u>p</u>aar (Tsp5) (Bernal *et al*, 2017; Velázquez *et al*, 2025). VgrG proteins form a trimeric complex at the distal end of the <u>H</u>aemolysin <u>c</u>oregulated <u>p</u>rotein (Hcp) tube and are essential for the assembly and function of the T6SS (Hachani *et al*, 2014). The last gene in the cluster, *tsp5*, is predicted to code for a PAAR (<u>p</u>roline-<u>a</u>lanine-<u>a</u>lanine-arginine repeat) protein. PAAR proteins assemble cone-shaped structures that sharpen and diversify the VgrG spike complex (Shneider *et al*, 2013).

Antiprokaryotic effector proteins delivered by the T6SS are often paired with cognate immunity proteins that protect the producing cell from self-intoxication, as well as interspecies (Basler *et al*, 2013) and intraspecies (George *et al*, 2024) competition. In this regard, previous work has demonstrated that Tki5 expression protects from Tke5-induced toxicity (Velázquez *et al*, 2025).

Tap3 is a protein belonging to the DUF4123 family, which has recently been identified as a group of T6SS adaptor proteins that play a crucial role in enabling the recognition and export of evolutionarily unrelated effectors (Colautti *et al*, 2024; Unterweger *et al*, 2015). DUF4123 proteins are often found downstream of *vgrG* or putative effector-encoding genes in predicted T6SS gene clusters. The presence of *vgrG3*, *tap3*, *tke5*, *tki5* and *tsp5* genes nearby within the K3-T6SS cluster underscores the complex and coordinated nature of the T6SS machinery, highlighting how structural components, adaptors, effectors, and immunity proteins work together to enable the system’s function. Although the relevance of the DUF4123 family of T6SS adaptor proteins, there is still little insight into the molecular mechanism they employ to tether their cognate effectors to the VgrG-PAAR spike complex. This is largely due to the limited structural information available. To provide insight into the complex interplay between these T6SS components, we have determined the 2.8 Å cryo-EM structure of Tke5 in complex with its cognate adaptor Tap3, revealing previously undescribed protein folds for Tap3 and Tke5. Our structure uncovers how Tap3 recognises the MIX domain of Tke5. Furthermore, we define the domain architecture of Tke5 and demonstrate that only an α-helical region located towards its C-terminal end is sufficient to kill intoxicated bacteria. In contrast, a β-rich region at the C-terminus is likely to enhance target membrane specificity. Together, our results provide a comprehensive framework for understanding how similar MIX-containing BTH_I2691 effectors might be recognised by their cognate Tap proteins, and we dissect their structural organisation and mechanism of action.

## RESULTS

### A 2.8 Å cryo-EM structure of the Tap3–Tke5 complex provides insight into Tke5’s molecular mechanisms for T6SS-loading and toxicity

The *tke5* gene is located within the K3-T6SS cluster of *Pseudomonas putida* KT2440, where it is found near the 3’ end along with genes coding for VgrG3, Tap3, its cognate immunity protein Tki5, and Tsp5 (**Fig. 1a**). Tap proteins co-evolved with MIX-containing effectors (Colautti *et al*, 2024). Interestingly, all characterised DUF4123 proteins function as adaptor proteins, providing a direct and essential physical connection between VgrG proteins and diverse families of cognate effectors that lack PAAR or “PAAR-like” N-terminal domains (Pei *et al*, 2020; Burkinshaw *et al*, 2018; Liang *et al*, 2015; Miyata *et al*, 2013; Unterweger *et al*, 2015; Colautti *et al*, 2024). Given the genomic context of the *tap3* and *tke5* genes and the function of other Tap proteins, we expected a direct interaction between Tap3 and Tke5 proteins. To demonstrate this, we carried out heterologous co-expression of Tap3 and Tke5 in *Escherichia coli*. Following co-expression, we are able to co-purify to homogeneity the Tap3–Tke5 complex in milligram quantities (**Fig. 1b**; see Methods for details), which affords solving its cryo-electron microscopy (cryo-EM) structure to 2.8 Å resolution (**Fig. 1c-d**, **Movie 1** and **Supplementary Fig. 1-2**; see Methods and **Supplementary Table 1** for details). The resulting cryo-EM density map shows high quality, permitting *de novo* atomic model building of the complex except for residues 1-12 (N-terminus) and 303-307 (C-terminus) of Tap3, and 1-16 (N-terminus), 267-273, 730-757, and 886-909 of Tke5.

**Fig. 1.**
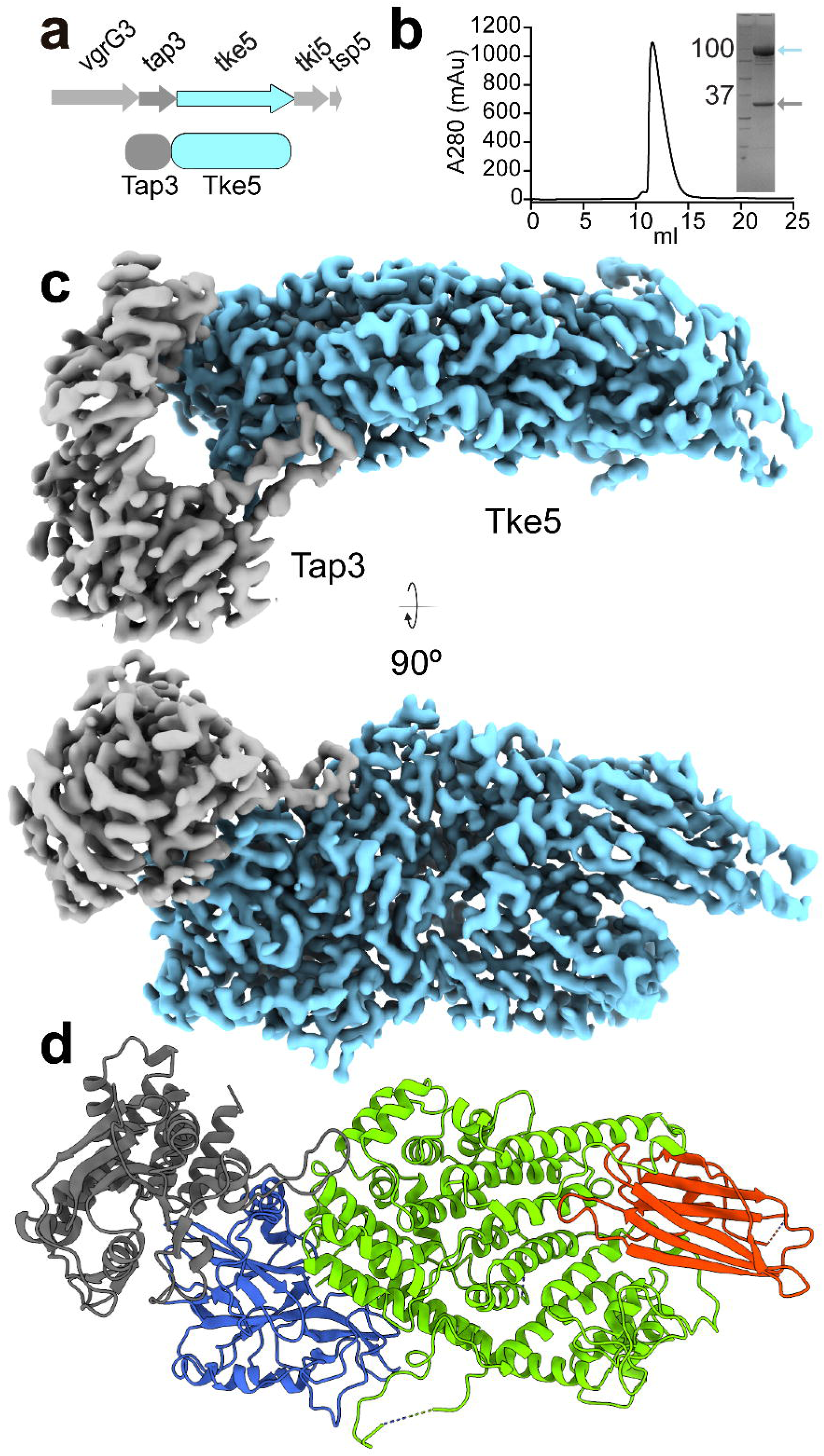
Genetic context, purification, and cryo-EM structure of the Tap3–Tke5 complex. **(a)** Schematic genetic organisation of the *vgrG3*, *tap3*, *tke5*, *tki5*, and *tsp5* genes. *tap3* and *tke5* are co-expressed from the pCOLADuet-1 plasmid, while *tki5* is expressed from the pSEVA624C plasmid to neutralise potential Tke5-induced toxicity. The arrows indicate the direction of transcription. **(b)** Gel filtration chromatogram of the purified Tap3–Tke5 complex. The complex eluted as a single, monodisperse peak, indicating a stable and homogeneous sample. The x-axis represents the elution volume (mL), and the y-axis represents the absorbance at 280 nm (mAU). The inset shows an SDS-PAGE analysis of the purified Tap3–Tke5 complex. The gel shows two distinct bands, corresponding to Tke5 and Tap3, confirming the presence of both proteins in the purified complex. This result is consistent with the mass spectrometry data (**Supplementary Data 1**). **(c)** Cryo-EM density of Tap3–Tke5 complex displayed perpendicular to the longest axis of the complex and rotated 90° clockwise (map post-processed with DeepEMhancer). **(d)** Cartoon representation of the Tap3–Tke5 atomic structure. Tap3 is shown in grey, and Tke5 is coloured in blue (MIX domain), green (α-helical region), and red (β-rich region).

Analysis of the Tap3–Tke5 complex reveals that Tap3 comprises two domains that assemble into a horseshoe-like fold (**Fig. 2a**), representing a novel fold without significant representatives in the Protein Databank (based on a Foldseek (Kim *et al*, 2025) search). The Tap3 N-terminal domain is an α/β-domain assembled by a central anti-parallel β-sheet containing six β-strands (shown in red) surrounded by a β-hairpin and six helices (three helices on each face of the β-sheet). The C-terminal domain is an α-helical bundle assembled by five helices (see **Supplementary Fig. 3** for the secondary structure arrangement).

**Fig. 2.**
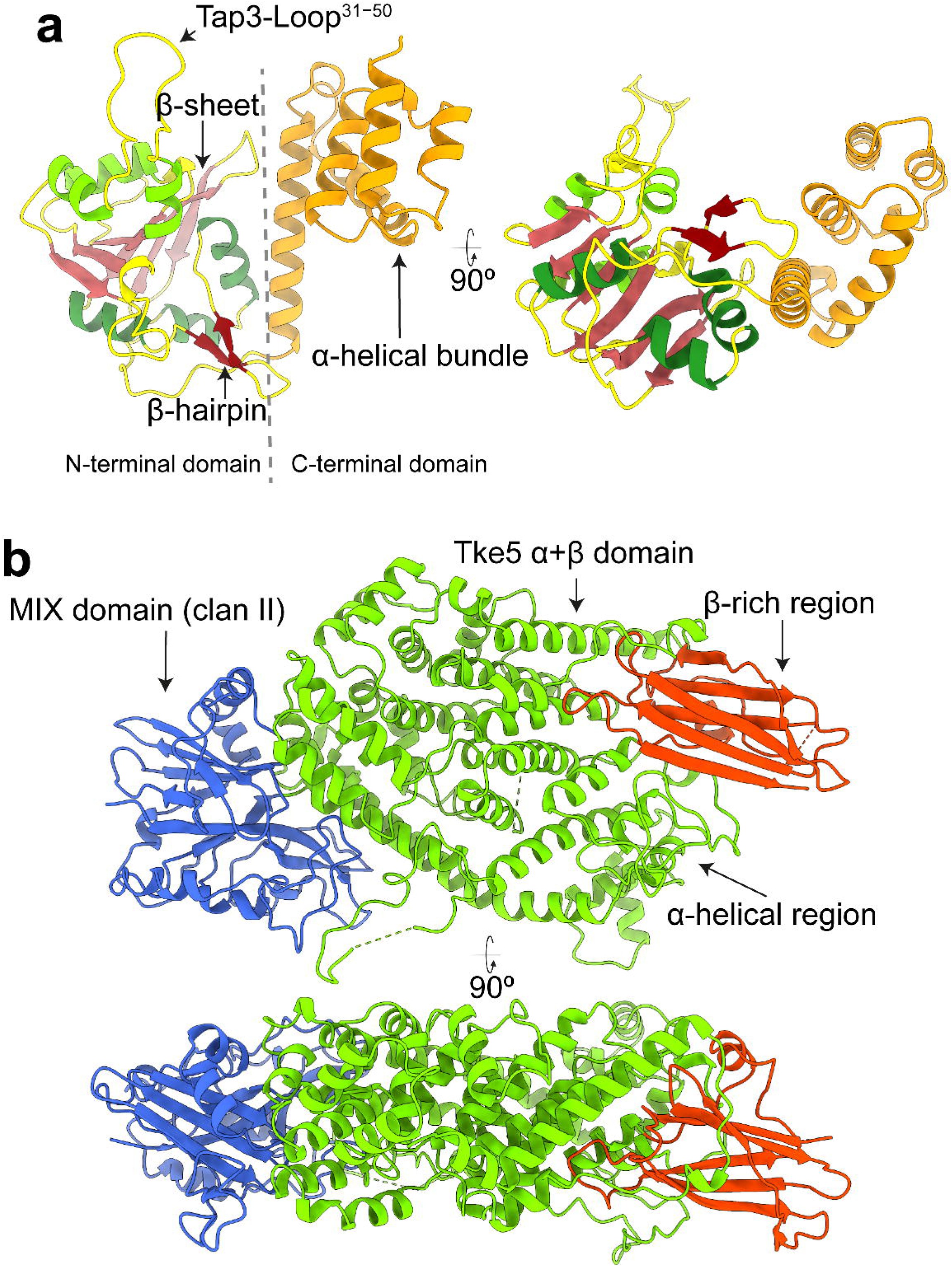
Overall domain architecture of Tap3 and Tke5. **(a)** Two 90° rotated views of Tap3 shown as cartoon representation, highlighting its N-terminal and C-terminal domains. The N-terminal domain (residues 1–198) adopts an α/β fold, featuring a central anti-parallel β-sheet (shown in red) flanked by a β-hairpin (shown in dark red) and six α-helices (three on each side of the β-sheet). The C-terminal domain (residues 199–302) forms an α-helical bundle comprising five helices (shown in orange). **(b)** Two views rotated 90° of Tke5 are shown as cartoon representations, highlighting its N-terminal and C-terminal domains. The N-terminal α/β domain contains a MIX fold (residues [17-238] shown in blue). The C-terminal α+β domain (residues [239-996]) is organised in an α-region (residues [239-863] shown in green) and a β-rich region (residues [864-996] shown in red). The β-rich region presents an immunoglobulin-like sandwich fold.

Tke5 is organised into two domains: an N-terminal α/β domain containing the MIX motif (residues [17-238]; shown in blue in **Fig. 2b**) and an α+β domain (residues [239-996]; shown in green and red). See **Supplementary Fig. 4, 5** for its secondary structure arrangement. The MIX motif-containing domain (hereafter named the MIX domain) mediates most of the binding with Tap3.

Remarkably, Tap3 includes a large loop (residues 31-50, named thereafter Tap3-Loop^31−50^) that protrudes outwards and accounts for ∼54% of the Tap3 residues that bind to Tke5 (**Fig. 3a-b**). Besides Tap3-Loop^31−50^, two more regions in Tap3 bind to Tke5, one of which is also located within the N-terminal domain (residues R75, E77, F78; **Fig. 3b**); and a second region within the C-terminal α-helical bundle (residues N251, L284, E286, S287, P288, Q289, A290, R293; **Fig. 3c**). In summary, the N-and C-terminal domains of Tap3 bind to the MIX domain of Tke5, except Tap3-Loop^31−50^ that extends its binding beyond the Tke5’s MIX domain and into Tke5’s α+β-domain. All hydrogen bonds between Tap3 and Tke5 are represented in **Supplementary Fig. 6** and **Supplementary Table 2**.

**Fig. 3.**
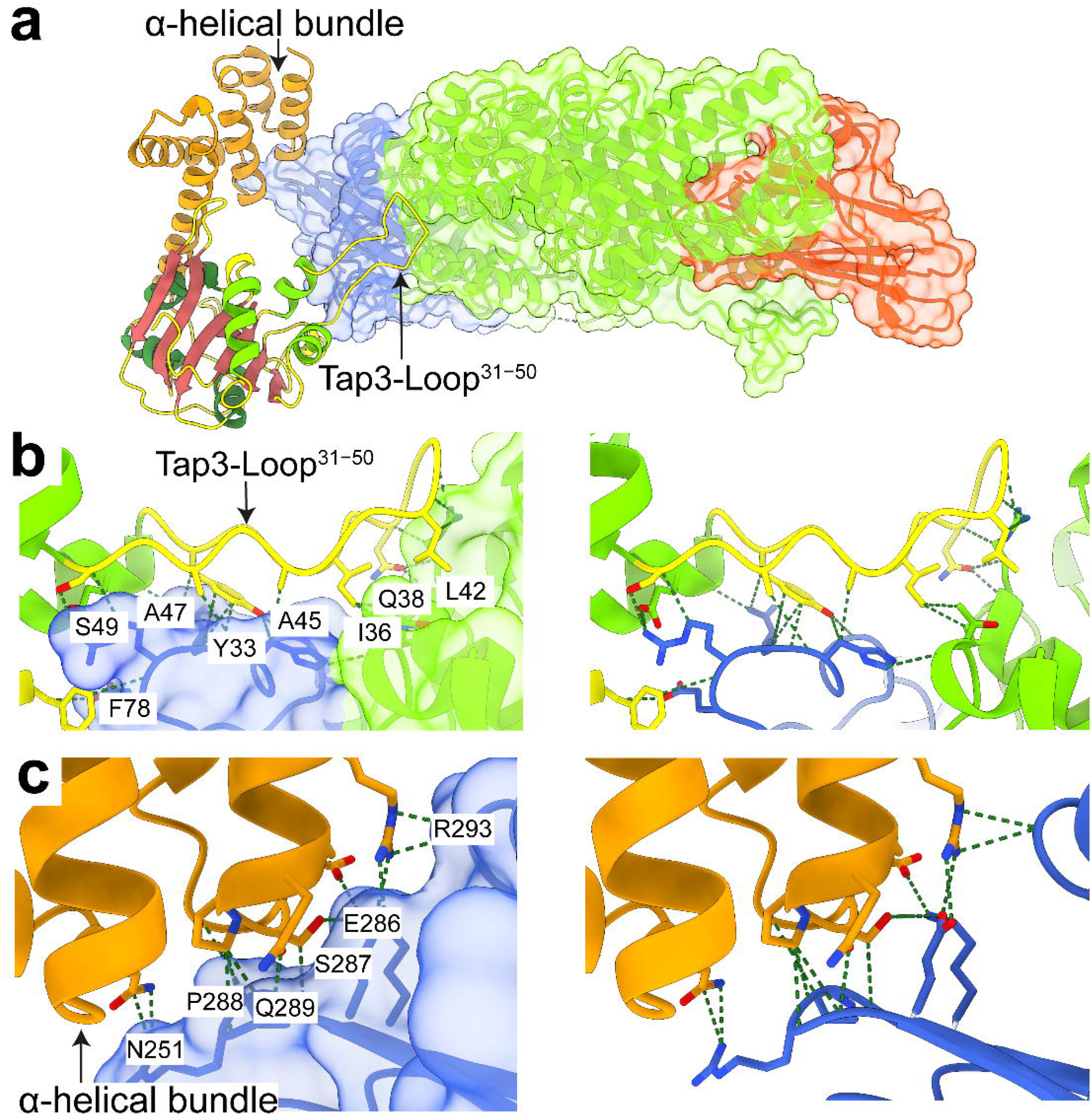
Structural insights into Tap3–Tke5 interactions. **(a)** Cartoon representation of the Tap3–Tke5 atomic structure. The solvent-excluded molecular surface of Tke5 is also shown. The colouring scheme is the same as for Fig. 2: the Tke5’s MIX fold is shown in blue, the α-region is shown in green, and the β-rich region is shown in red. Tap3’s N-terminal domain is colour-coded by secondary structure elements, and the C-terminal α-helical domain is shown in orange. **(b)** Close-up view of the prominent loop (Tap3-Loop^31–50^) extending from the N-terminal domain of Tap3 and interacting with Tke5. This loop protrudes outward and accounts for 54% of the Tap3 residues contacting Tke5. The solvent-excluded molecular surface of Tke5 is shown on the left panel and is hidden on the right panel. **(c)** Close-up view showing the interactions between Tap3’s C-terminal α-helical domain and Tke5’s MIX fold. The solvent-excluded molecular surface of Tke5 is shown on the left panel and is hidden on the right panel.

### The MIX fold is a conserved structural scaffold for effector specificity and adaptor recruitment

The MIX motif is commonly found in antibacterial T6SS effectors, and sequence clustering analyses have identified up to five distinct clans (MIX I to MIX V) (Salomon *et al*, 2014). This MIX motif was defined by a conserved central sequence, hRxGhhYhh (where *h* denotes a hydrophobic residue), flanked by less conserved segments at the N-terminus (shhPhR) and the C-terminus (hhF/YSxxxWS/T) (Salomon *et al*, 2014). The cryo-EM structure of Tke5 reveals that the MIX fold has no close structural homologs in the Protein Data Bank (as determined by Foldseek (Kim *et al*, 2025) analysis). This fold features a distinctive pyramid-like architecture composed of two central β-sheets connected by variable-length loops and six α-helices (**Fig. 4**; **Supplementary Fig. 7**).

**Fig. 4.**
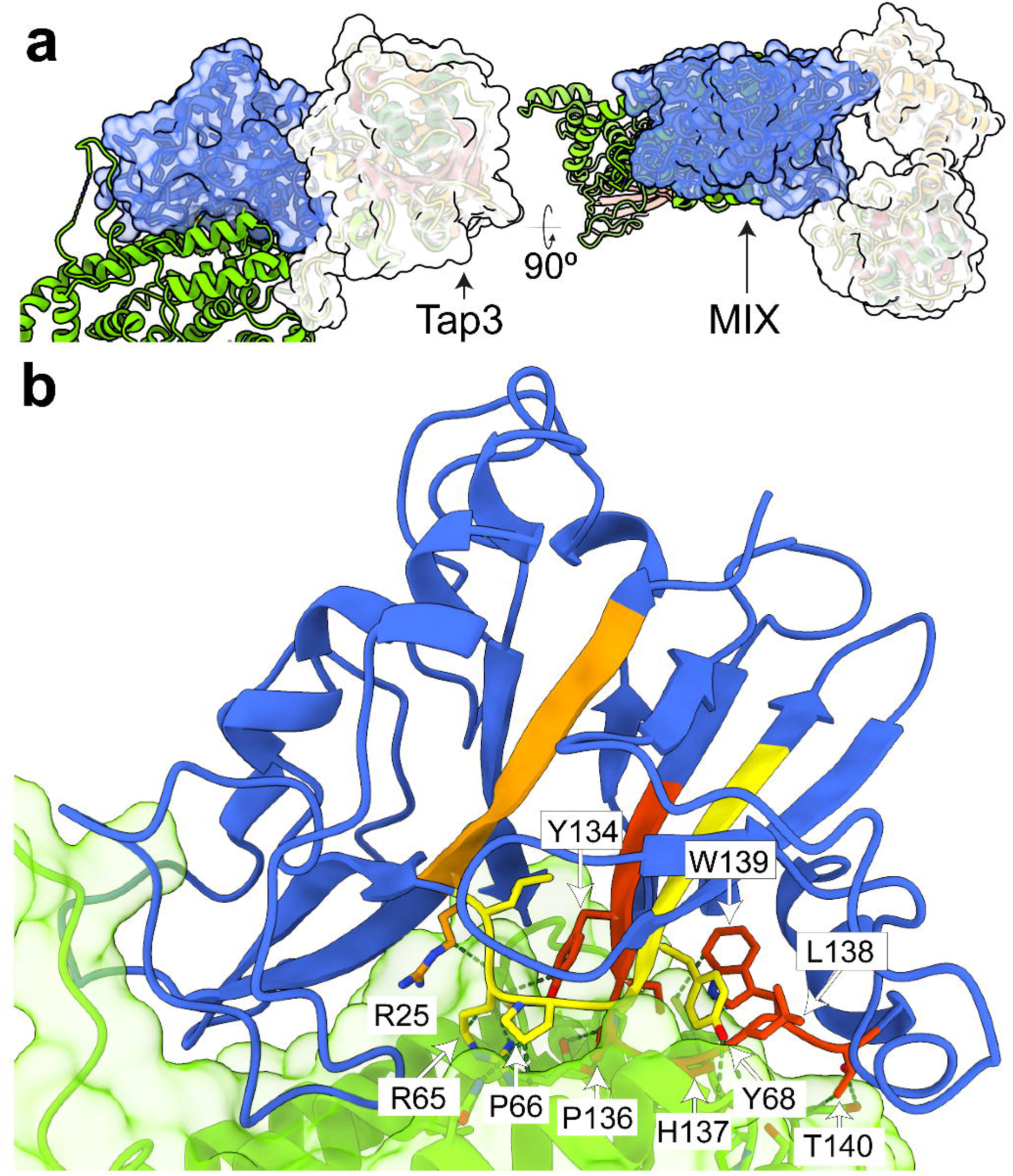
The MIX motif and its role in Tke5 domain specificity. **(a)** Two 90° anti-clockwise rotated views of the Tap3–Tke5 atomic structure. The solvent-excluded molecular surface of the MIX domain and Tap3 are also shown. The colouring scheme is the same as for Fig. 2 and 3. **(b)** Close-up view of the MIX–α+β domain interface, highlighting the MIX motif with residues 20–25 (PILPVR) in orange, 64–72 (LRPGYVYVF) in yellow, and 132–140 (IGYSPHLWT) in red. Key interfacial contacts are indicated, including salt bridges formed by R25 and R65, hydrogen bonds involving Y68, L138, and T140, and additional contacts contributed by P66, G67, Y68, Y134, P136, H137, L138, W139, and T140.

The specific sequence of the MIX motif in Tke5 comprises residues ^20^PIL<u>P</u>V<u>R</u>^25^, ^64^L<u>R</u>P<u>G</u>YV<u>Y</u>VF^72^, and ^132^IG<u>YS</u>PHLW<u>T</u>^140^. R25 and R65 residues are forming salt bridges with the α+β-domain; Y68, L138, and T140 are involved in hydrogen bonds with the α+β-domain; and P66, G67, Y68, Y134, P136, H137, L138, W139, and T140 are also in contact with the α+β-domain (**Fig. 4b**). Thus, the structure reveals that most of the residues in the MIX motif localise at the interface between MIX and α+β-domains, which suggests that MIX domains might provide specificity to their toxic domains found in MIX-containing T6SS effectors, as well for recognising their corresponding cognate Tap proteins. This latter function has been recently demonstrated for two *Pseudomonas aeruginosa* adaptor proteins, Tap6 and Tap14, which specifically tether unrelated T6SS effectors to their respective VgrG partners (Colautti *et al*, 2024). All hydrogen bonds between the MIX domain and the α+β domain are represented in **Supplementary Fig. 8** and **Supplementary Table 3**.

### The α-helical region of Tke5 is essential for toxicity and immunity-mediated neutralisation

The toxic activity encoded within Tke5 is responsible for bacterial killing, membrane depolarisation, and pore formation (Velázquez *et al*, 2025). Based on Tke5’s domain architecture, the pore-forming activity is likely mediated by its C-terminal α+β domain. This domain can be subdivided into two distinct regions: an N-terminal α-helical region (residues 239–863 shown in grey and green; **Fig. 5a**) and a C-terminal β-rich region (residues 864–996 shown in red; **Fig. 5a**). The α-helical region is assembled by 34 helices (∼ 62% of residues). These helices are closely packed, forming a compact structure. Prediction of transmembrane helices based on primary sequence suggests that five segments within the α-helical region might have a propensity for membrane insertion (predicted by TMHMM 2.0 (Krogh *et al*, 2001); **Supplementary Fig. 9**). These five segments are contained within residues 608-813 and include all or part of 11 helices. Six of these helices are buried in the core of the α-helical region, while the remaining five are located on the surface (residues 608-813 shown in green; **Fig. 5a**; **Supplementary Table 4**). Given that this structure is most likely representing a pre-pore state of Tke5 found in solution, one would expect major conformational changes upon membrane insertion, which might drive some of these helices to insert into the hydrophobic core of the target membrane, resulting in the assembly of a pore-state.

**Fig. 5.**
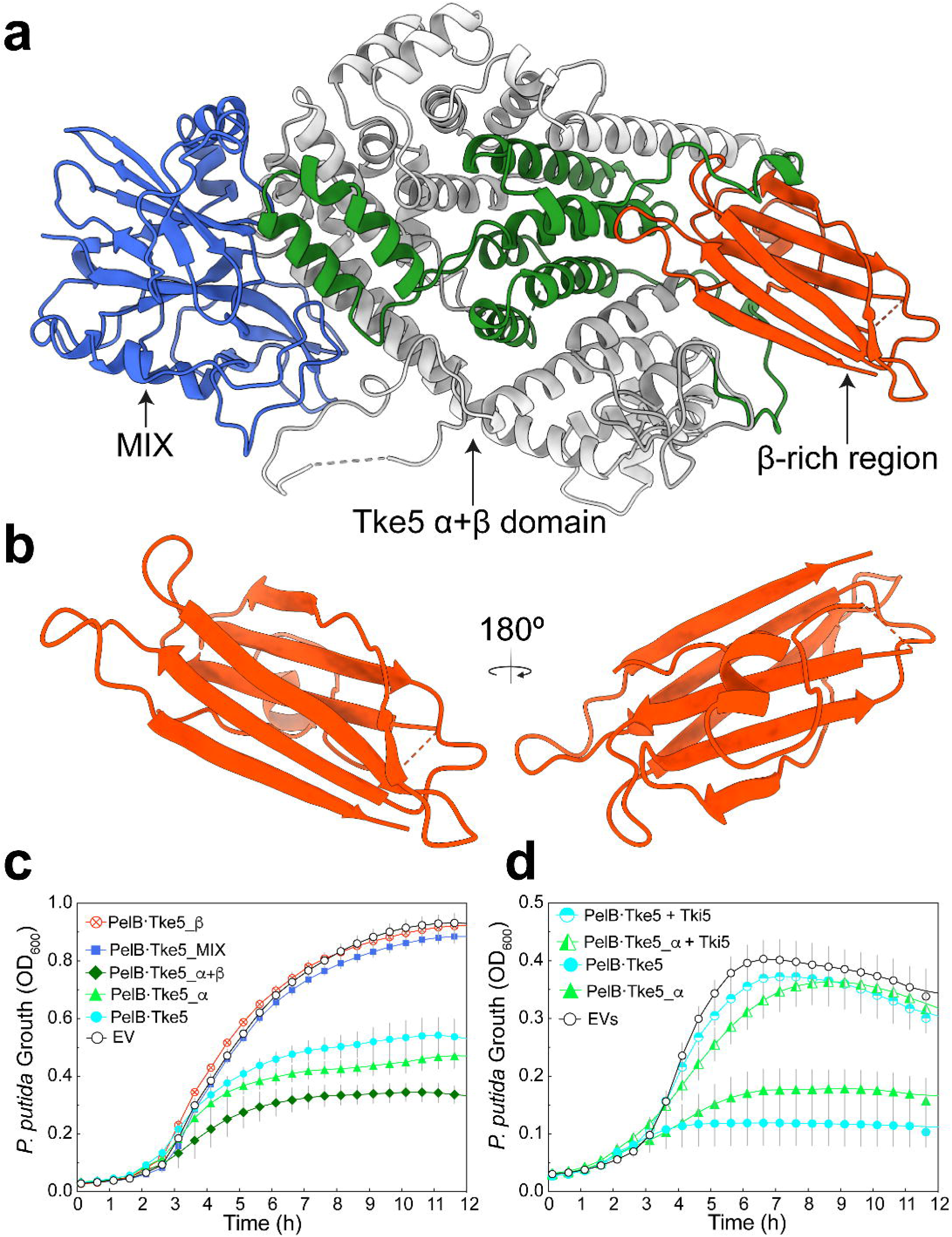
The α-helical region of Tke5 mediates toxicity and is neutralised by cognate immunity protein Tki5. **(a)** Domain architecture of Tke5 showing the N-terminal MIX domain (residues 17–238, blue), α-helical pore-forming region (residues 239–863, grey and green), and C-terminal β-rich region (residues 864–996, green). Predicted transmembrane helices (green) are mapped onto the α-helical region, which correspond to residues 622-644 (FLGLLTSGGGGGLNAGMLWFNIL), 663-685 (LGFASSVFGVIGAAAATLVSV-RA), 695-717 (ISTAPGMAFGNGIINFLTSNLFA), 744-766 (TALGSILVL) and 781-803 (FAGWTAAGIALIGATLIGGGLFL). **(b)** Two 180° rotated views of the β-rich region that presents an immunoglobulin-like sandwich fold located at the C-terminal of Tke5. **(c)** Toxicity assays of Tke5 and truncation constructs in *P. putida*. Full-length Tke5 (PelB·Tke5), the α+β domain (PelB·Tke5_α+β) and the α-helical region alone (PelB·Tke5_α) show comparable toxicity. The MIX domain (PelB·Tke5_MIX) and β-rich region (PelB·Tke5_β) are non-toxic. **(c)** Neutralisation of Tke5 toxicity by its cognate immunity protein Tki5. Co-expression of Tki5 rescues cell growth inhibited by PelB·Tke5, or PelB·Tke5_α, confirming the specificity of Tki5 for the α-helical toxic domain.

The C-terminal β-rich region presents an immunoglobulin-like sandwich fold. The sandwich is assembled by two antiparallel β-sheets, one with three strands and the other with four. One face of the sandwich is packed against the only helix in this domain (residues 864-996 shown in red; **Fig. 5a-b**). Tke5 homologues are found across the Proteobacteria (Pseudomonadota) phylum, where they are predominantly annotated as T6SS effector BTH_I2691 family proteins. Interestingly, some family members do not contain the C-terminal β-rich region, which led us to hypothesise that the α-helical region might be sufficient for toxicity.

To test our hypothesis, we measured the toxicity of Tke5 full-length and deletion constructs when expressed from plasmid pS238D•M in *P. putida* KT2440 (**Fig. 5c**). Tke5 full-length and deletion constructs contain an N-terminal PelB leader sequence for targeting the Tke5 constructs to the secretory (Sec) pathway. We engineer the following constructs: the PelB−containing Tke5 full length (PelB·Tke5) and deletion constructs PelB·Tke5^1-238^ (PelB·Tke5_MIX), PelB·Tke5^239-996^ (PelB·Tke5_α+β), PelB·Tke5^239-863^ (PelB·Tke5_α), PelB·Tke5^864-996^ (PelB·Tke5_β). As expected, PelB·Tke5 and PelB·Tke5_α+β constructs are toxic, with comparable toxicity levels, indicating that the MIX domain is not necessary for toxicity. In line with this result, the PelB·Tke5_MIX construct is non-toxic. Importantly, the PelB·Tke5_α construct that lacks the MIX and the β-rich region is also toxic, indicating that the α-helical region is sufficient for toxicity and that the β-rich region might play a secondary role in Tke5’s activity.

To determine that the toxicity observed for PelB·Tke5 and PelB·Tke5_α directly depends on their activity, we co-expressed these constructs with the Tke5 cognate immunity protein Tki5 encoded on a pSEVA624C plasmid (see Methods for details). As expected, the expression of Tki5 protects against the toxicity of PelB·Tke5 and PelB·Tke5_α (**Fig. 5d**), further supporting the idea that the toxic domain resides within the α-region and that the immunity protein neutralises toxicity by specifically targeting it.

It has previously been demonstrated that Tke5 is a bactericidal antimicrobial toxin that induces cell depolarisation (Velázquez *et al*, 2025). Our findings now establish that its bactericidal activity is directly mediated by the α-helical region of Tke5, which alone suffices to kill intoxicated bacteria. This suggests that while the α-helical region encodes the core pore-forming toxin responsible for disrupting ion homeostasis, the β-rich C-terminal region might function as a receptor-binding module to enhance target membrane specificity or effector stability. Crucially, the Tki5 neutralisation of the toxicity induced by PelB·Tke5_α confirms that the immunity protein directly counteracts the effector’s toxic activity rather than interfering with other domains or membrane recruitment mechanisms.

### Proposed Tke5’s pore-forming molecular mechanism

The comprehensive analysis of Tke5’s structure and the functional dissection of each domain allows for the proposal of a molecular mechanism for its pore-forming activity. Tke5 might operate through a sophisticated, multi-step process, culminating, as previously demonstrated, in the disruption of target bacterial membrane potential without causing widespread cellular lysis (Velázquez *et al*, 2025).

The mechanism begins with the delivery of Tke5 into the periplasmic space of the target bacterium. Once in the periplasm, Tke5 must interact with the inner membrane to exert its toxic effect. Our model hypothesises that the C-terminal β-rich region (residues 864-996), which adopts an immunoglobulin-like sandwich fold, functions as a receptor-binding domain (RBD). This hypothesis is based on the fact that the β-rich region is not essential for toxicity (**Fig. 5c**) and the predicted position of Tke5 on a model Gram-negative inner membrane by the PPM 3.0 server (Protein Property Prediction and Modelling Server (Lomize *et al*, 2022); see Methods for details). PPM 3.0 predicts that the pre-pore state of Tke5 binds to the membrane via the RBD domain (**Fig. 6a**). In particular, this binding is predicted to occur through the most distal end of the RBD, including a disordered loop (residues 886-909) not visible in the cryo-EM map that we modelled using AlphaFold_unmasked (Mirabello *et al*, 2024). The predicted interaction occurs with one of the leaflets of the membrane, suggesting an initial, perhaps low-affinity, interaction with the lipid bilayer. This membrane-sensing role of the RBD could be relevant for guiding Tke5 to its target membrane and potentially orienting it for subsequent insertion.

**Fig. 6.**
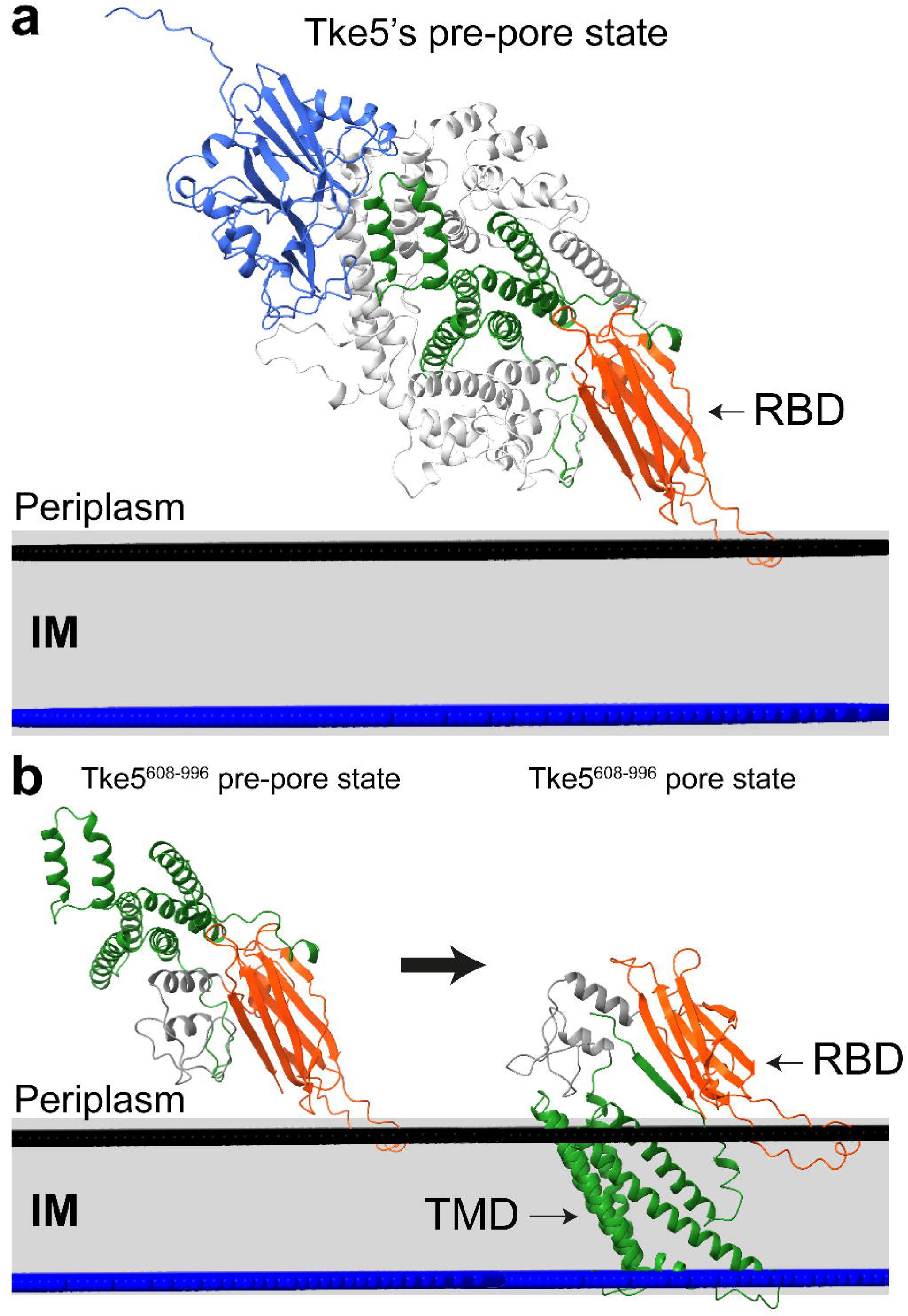
Proposed molecular mechanism for Tke5 pore formation. **(a)** Model of Tke5 interaction with the inner membrane of a target bacterium. Following delivery into the periplasmic space via the T6SS, Tke5 is hypothesised to initially interact with the inner membrane through its C-terminal β-rich receptor-binding domain (RBD, residues 864-996, shown in red). This initial binding, predicted by PPM 3.0, occurs via the distal end of the RBD, including a modelled disordered loop (residues 886-909). In its soluble, pre-pore state, the α-helical region (residues 239-863, shown in grey and green) contains five segments (residues 608-813, shown in green) with a propensity for membrane insertion. **(b)** Conformational change of Tke5 upon membrane interaction. A comparison between the AlphaFold3 (AF3) predicted structure for Tke5 (residues 608-996) and the corresponding region in the cryo-EM structure suggests that predicted transmembrane (TM) helices in Tke5 undergo an extensive conformational change upon membrane interaction. Combined with PPM 3.0 calculations, these predictions suggest that TM helices cross the lipid bilayer, forming a Transmembrane Domain (TMD) that is proposed to be responsible for pore formation upon membrane binding. This multi-step process results in the disruption of the target bacterial membrane potential. PPM 3.0 calculations are performed with a model *E. coli* inner membrane (PE:PG:CL (79:19:2)). The membrane is shown schematically in a perpendicular orientation, with black and blue surfaces representing the interface between the lipid polar head groups and the acyl chains.

Following this initial membrane sensing by the RBD, the α-helical region (residues 239-863) of Tke5, which contains five segments with a strong propensity for membrane insertion, might undergo significant conformational changes as predicted by AlphaFold 3 (AF3) (Abramson *et al*, 2024) (residues 608-813; predicted transmembrane domain (TMD) shown in dark green in **Fig. 6**). In its soluble, pre-pore state, many of these helices are buried within the protein’s core or located on its surface (**Fig. 6a**). However, comparison between the AF3 predicted structure for Tke5 residues 608-996 (Tke5^608-996^) and the corresponding region in the cryo-EM structure suggests that the predicted TMD in Tke5 undergoes an extensive conformational change upon membrane interaction (**Fig. 6b**). The AF3 prediction of Tke5^608-996^, combined with the PPM 3.0 calculations, suggest that the predicted TMD crosses the lipid bilayer. Taken together, these predictions suggest that the TMD is most likely responsible for pore formation.

## DISCUSSION

### Structural insights into Tap-effector recognition

Our 2.8 Å cryo-EM structure of the Tap3-Tke5 complex from *Pseudomonas putida* KT2440 provides the first atomic-level insight into how DUF4123 adaptor proteins recognise their cognate effectors. This high-resolution structure reveals novel protein folds for both Tap3 and Tke5. Tap3 adopts a unique horseshoe-like bilobed architecture, while Tke5 is organised into an N-terminal α/β domain containing the MIX motif (Clan II) and a C-terminal α+β domain. Crucially, the Tke5 MIX domain mediates the majority of the binding with Tap3. A remarkable feature of this interaction is the prominent Tap3-Loop^31-50^, which protrudes from Tap3’s N-terminal domain and accounts for approximately 54% of the Tap3 residues involved in binding to Tke5. This loop extends its interaction beyond Tke5’s canonical MIX domain into its α+β-domain, indicating a broad and intimate interface.

Previous research had demonstrated a direct interaction between another Tap protein (Tap6) and its cognate MIX-containing BTH_I2691 effector (Ptx2). The work combined AlphaFold 3 (AF3) predictions with extensive experimental validation (Colautti *et al*, 2024). While AF3 confidently predicted Tap6’s bilobed architecture and its N-terminal lobe interacting with a C-terminal helix-turn-helix (HTH) motif found in its cognate VgrG6, it was notably unable to confidently predict the structure of Ptx2 or the complex formed between Ptx2 and Tap6 due to Ptx2 having few close sequence or structural homologs (Colautti *et al*, 2024). Despite this, the Tap6-Ptx2 binding mechanism was strongly supported by experimental evidence, including bacterial competition assays that demonstrated Tap6’s essential role in Ptx2 export, *in vitro* size exclusion chromatography showing stable complex formation, and *in vivo* co-immunoprecipitation, which confirmed that Tap6 mediates the interaction between Ptx2 and VgrG6. Based on the general bilobed architecture of DUF4123 proteins, it was proposed that Tap6’s variable C-terminal lobe is responsible for recognising and binding Ptx2 (Colautti *et al*, 2024).

The sequence homology between Tap3 and Tap6 is strikingly low, at only ∼ 16% (**Supplementary Fig. 10a**). Similarly, their corresponding cognate effectors, Tke5 and Ptx2, despite containing MIX domains, also share very low sequence identity (∼ 17%; **Supplementary Fig. 10b**). The presence of Tap3-Loop^31-50^, which is not present in Tap6 and accounts for over half of Tap3’s binding residues to Tke5, marks a clear structural and mechanistic divergence in effector recognition. This divergence in these two Tap proteins likely reflects differences in how they specifically recognise their cognate effectors. Although both Tap3 and Tap6 recognise MIX-containing BTH_I2691 effectors, the distinct classification of Tke5 containing a Clan II MIX domain and Ptx2 containing a Clan I MIX domain further underscores the need for specialised adaptor recognition. This remarkable evolutionary plasticity suggests DUF4123 adaptors employ diverse structural strategies to specifically recognise a broad repertoire of effectors, thereby expanding the functional reach of the T6SS.

### Electrophysiological properties and diverse mechanisms of T6SS Pore-Forming Toxins causing bacterial cell depolarization

The T6SS deploys a diverse arsenal of toxic effector proteins, including PFTs such as VasX (Miyata *et al*, 2013), Tme1 (Fridman *et al*, 2020), Tme2 (Fridman *et al*, 2020), Ptx2 (Colautti *et al*, 2024), Tse5 (Rojas-Palomino *et al*, 2025a; González-Magaña *et al*, 2022), Tse4 (Rojas-Palomino *et al*, 2025b), Ssp4 (Reglinski *et al*, 2024), and Ssp6 (Mariano *et al*, 2019). The unifying principle among these PFTs is their ability to induce membrane depolarisation as a primary mechanism of bacterial intoxication. The electrophysiologic properties of Tse5 (Rojas-Palomino *et al*, 2025a; González-Magaña *et al*, 2022), Tse4 (Rojas-Palomino *et al*, 2025b), Ssp4 (Reglinski *et al*, 2024), and Ssp6 (Mariano *et al*, 2019), as well as Tke5 (Velázquez *et al*, 2025), have been studied in detail, demonstrating that their toxicity is due to their capacity to assemble ion-selective pores in the inner membrane of Gram-negative bacteria, generally preferring cations over anions. Nonetheless, their specific biophysical properties and modes of action exhibit remarkable diversity (**Table 1**).

**Table 1.** Comparative electrophysiological analysis of Tke5, Tse4, Tse5, Ssp4, and Ssp6.

| Characteristic | **Tke5** (Velázquez *et al*, 2025) | **Tse4** (Rojas-Palomino *et al*, 2025b) | **Tse5** (González-Magaña *et al*, 2022; Rojas-Palomino *et al*, 2025a) | **Ssp4** (Reglinski *et al*, 2024) | **Ssp6** (Mariano *et al*, 2019) |
| --- | --- | --- | --- | --- | --- |
| Source Organism | *P. putida* KT2440 | *P. aeruginosa* PAO1 | *P. aeruginosa* PAO1 | *Serratia marcescens* Db10 | *Serratia marcescens* Db10 |
| Toxin Family/Type | BTH_I2691 family PFT | T6SS effector PFT | Rhs toxin, T6SS PFT | Ssp4-like proteins | * Tpe1 |
| Primary Mechanism of Action | Membrane Depolarization | Membrane Depolarization | Membrane Depolarization | Membrane Depolarization | Membrane Depolarization |
| Membrane Damage | No large damage | No large damage | No large damage | No large damage | No large damage |
| Pore Conductance | € 33 ± 4 pS | € 20 ± 3 pS | € 196 ± 8 pS ‡ | § 18.4 ± 0.64 pS | - |
| Estimated Pore Radius (nm) | ~0.4 | ≤ 0.4 | ~ 0.9 | - | - |
| Reversal Potential (mV) | @ -31 ± 2 | @ -20 (variable) | @ -14.0 ± 2.1 | \$ -14.8 ± 1.6 | \$ -55.8 ± 3.4 |
| Ion permeability ratio (P<sub>K+</sub>/P<sub>Cl-</sub>) | 10 ± 3 | 4.0 ± 1.9 | 3.8 ± 0.9 | - | - |
| Ion Selectivity | Cation-selective | Cation-selective | Cation-selective | Cation-selective | Cation-selective |
| Specific cation preference | - | Na<sup>+</sup>, Li<sup>+</sup> > K<sup>+</sup> | no cation specificity | no cation specificity | # K<sup>+</sup> ≈ Na<sup>+</sup> |
| Voltage Dependence / Rectification | Ohmic | Ohmic (NaCl, KCl) & Outward Rectifying (KCl) | Initial voltage-dependent, then Ohmic | Ohmic | - |
| Proposed Insertion Mechanism | Periplasmic insertion | Periplasmic insertion | Periplasmic insertion | Periplasmic insertion | Periplasmic insertion |
| Immunity Protein | Tki5 | Tsi4 | Tsi5 | Sip4 | Sip6 |

\* T6SS-dependent pore-forming effector € Measured in symmetrical 150/150 mM KCl (cis/trans chambers) and lipid membrane assembled from *E. coli* polar lipid extract § Measured in symmetrical 510 mM KCl and a membrane assembled from bovine phosphatidylethanolamine lipids. ‡ Unpublished data @ Measured in asymmetrical 250/50 mM KCl (cis/trans chambers) and lipid membrane assembled from *E. coli* polar lipid extract \$ Measured in asymmetrical 500/210 mM KCl (cis/trans chambers) and lipid membrane assembled from bovine phosphatidylethanolamine lipid

Tke5 forms subnanometric (∼ 0.4 nm radius) ohmic pores with a strong preference for cations over anions (PK^+^/PCl^-^ = 10 ± 3), primarily causing depolarisation via Na^+^ influx (Velázquez *et al*, 2025). Tse4, which also forms narrow pores (with a radius of ≤ 0.4 nm), uniquely combines ohmic and diode-like rectifying channels.(Rojas-Palomino *et al*, 2025b) Thus, Tse4 exhibits an exquisite mechanism of action whereby initial depolarisation is postulated to result from the influx of Na^+^ through Tse4-assembled Ohmic pores that operate under a resting membrane potential (Rojas-Palomino *et al*, 2025b). These pores show mild cation selectivity for Na^+^ and Li^+^ over K^+^ (P ^+^/PCl^-^ = 4.0 ± 1.9). After cell depolarisation (i.e., at positive membrane potentials), Tse4-rectifying pores activate to facilitate K^+^ efflux, thereby balancing charge without altering pH (LaCourse *et al*, 2018). Tse5 forms nanometric proteolipidic pores (∼ 0.9 nm radius) that initially exhibit voltage-dependent conductance before stabilising to ohmic behaviour (González-Magaña *et al*, 2022; Rojas-Palomino *et al*, 2025a). Its weak cation selectivity (P ^+^/PCl^-^ = 3.81 ± 0.86) is significantly influenced by membrane lipid composition (Rojas-Palomino *et al*, 2025a).

Ssp4 forms cation-selective pores (18.4 ± 0.64 pS conductance) that also showed cation selectivity. Notably, Ssp4 intoxication leads to an increase in intracellular reactive oxygen species (ROS) (Reglinski *et al*, 2024). Ssp6, another cation-selective PFT, shows a strong preference for monovalent cations (Na^+^, K^+^) over divalent ones (Ca^2+^) and is not permeable to protons. Notably, Ssp6 also increases the permeability of the outer membrane, distinct from its effect on inner membrane depolarisation (Mariano *et al*, 2019).

Despite their shared outcome of membrane depolarisation and non-lytic growth inhibition, these subtle differences in biophysical characteristics, such as voltage-dependent rectification and specific cation selectivity, highlight the evolutionary fine-tuning of these PFTs to achieve specific physiological outcomes within the target cell. Tse4’s rectifying behaviour may allow for a more controlled K^+^ efflux, potentially synergising with other effectors or adapting to specific intracellular conditions. At the same time, Tke5’s mechanism appears more straightforward in its cation influx-driven depolarisation. Tse5’s unique encapsulation and release mechanism, along with its strong lipid dependence, further illustrate the diverse strategies employed by these toxins. This convergence on the inner membrane as a target underscores its fundamental importance in bacterial physiology, suggesting that PFTs that specifically and efficiently disrupt its electrochemical potential represent a highly successful evolutionary strategy for interbacterial competition.

### Summary

Our study elucidates the structural and functional basis of *Pseudomonas putida* Tke5, a pore-forming toxin that antagonises plant pathogens. The cryo-EM structure of the Tap3– Tke5 complex at 2.8 Å resolution reveals two novel protein folds, providing the first atomic-level insight into how MIX motifs mediate effector-adaptor interactions. We demonstrate that Tap3, a DUF4123 family adaptor, specifically recognises Tke5. Functional dissection identifies the α-helical region of Tke5 as the core pore-forming module responsible for membrane depolarisation and bactericidal activity. While the C-terminal β-rich region likely enhances target membrane specificity. Crucially, the immunity protein Tki5 directly neutralises the α-helical toxicity, underscoring the evolutionary precision of effector-immunity pairing. These findings establish a model for MIX-dependent effector recruitment, emphasising the modular architecture of T6SS toxins. By integrating structural and functional analyses, this work advances our understanding of bacterial competition mechanisms and provides significant insight into the widespread BTH_I2691 protein family.

## METHODS

### Construction of plasmids and bacterial strains

For a detailed overview of all strains and plasmids used in this study, please refer to **Supplementary Data 1**. All cloning procedures were conducted as described in previous work (Velázquez *et al*, 2025). In brief, the gene coding for Tke5 (PP2612) was amplified from genomic DNA isolated from the *Pseudomonas putida* KT2440 strain using P1-P2 primers. Additionally, P2-P3 primers were employed to amplify Tke5 and fuse a PelB signal sequence at the 5’ end (**Supplementary Data 1**). The *tke5* and *pelB·tke5* genes were first cloned into pJET1.2/blunt (Thermo Scientific™) and verified by Sanger sequencing using Macrogen Inc. services. Subsequently, *tke5* and *pelB·tke5* were subcloned into the pS238D•M vector at NheI/BamHI sites, resulting in pS238D•*tke5* and pS238D•*pelB·tke5,* respectively (**Supplementary Data 1**). This broad host range and medium copy number vector originates from the SEVA collection (Martínez-García *et al*, 2023a).

Then, pS238D•*pelB·tke5* was used as the parental vector to generate a series of plasmids containing specific Tke5 domains fused to the N-terminal signal sequence. Tke5_β (864-996) was amplified using primers P4-P5 and then subcloned to generate pS238D•*pelB·tke5_β.* The Tke5-MIX domain (1-238) was amplified using primers P6 and P7 and then subcloned to generate pS238D•*pelB·tke5_MIX.* The coding DNA region for Tke5_α*+β* (239-996) was amplified using primers P8 and P5 and then subcloned to generate pS238D•*pelB·tke5_α+β.* Tke5_α (239-863) was amplified using primers P8-P9 and then subcloned to generate pS238D•*pelB·tke5_α.* (**Supplementary Data 1**). Colony PCR using primers P10-P11 were used to screen colonies for pS238D•M derivatives and their subsequent sequencing.

Cloning of the *tki5* (PP2611) gene was performed as described in previous work (Velázquez *et al*, 2025). In brief, the *tki5* (PP2611) gene with an artificial RBS region (TTTAAAGGAGATATACAA) at the 5’ end was synthesised and cloned between the KpnI and HindIII restriction sites into pSEVA424 vector by GenScript (**Supplementary Data 1**). Then, both the RBS and *tki5* were subcloned from pSEVA424•*tki5* into pSEVA624C at KpnI/HindIII sites to generate pSEVA624C•*tki5* (**Supplementary Data 1**). This vector is a derivative of pSEVA621 that comes from the SEVA collection (Silva-Rocha *et al*, 2013; Martínez-García *et al*, 2023b) and was generated in a previous work where *lacI*^q^-*Ptrc* cargo from pSEVA234C (Nikel *et al*, 2022) was inserted in the MCS using PacI/HindIII (Velázquez *et al*, 2025). By colony PCR using P12-P13 primers, colonies harbouring the insert *tki5* were screened and sequenced.

The Tke5 coding sequence was subcloned from pS238D•*tke5* into the multiple cloning site 2 (*mcs-2*) of pCOLADuet^TM^-1. This subcloning was performed by GenScript (GenScript, New Jersey, USA), resulting in the plasmid pCOLADuet-1•*tke5*. The *tap3* (PP2613) gene was synthesised and cloned into the *mcs-1* of pCOLADuet-1•*tke5* by using the GenScript service. This subcloning conserves the 5’ 6xHis-tag of the *mcs-1* and leads to pCOLADuet-1•6x*his-tap3*-*tke5* (**Supplementary Data 1**).

### Expression, purification and cryo-EM vitrification of Tap3–Tke5 complex

*Escherichia coli* Bl21(DE3) cells co-transformed with pCOLADuet-1•6xhis-*tap3*-*tke5* and pSEVA624C•*tki5* plasmids were grown overnight at 37 °C with constant agitation in 200 mL of LB media containing 50 μg/mL kanamycin (Km), 20 μg/mL gentamycin (Gm) and 0.2% glucose. For Tap3 and Tke5 overexpression, bacterial cultures were diluted to an initial OD_600_ of 0.1 in flasks with 2 L of fresh LB medium supplemented with Km and Gm at the same concentrations but lacking glucose, and let them grow at 37 °C with shaking conditions. In the absence of glucose, Tki5 is expressed to neutralise possible Tke5-induced toxicity. When cells reached an OD_600_ value of 0.7, protein expression was induced by adding isopropyl β-D-1-thiogalactopyranoside (IPTG) at a final concentration of 1 mM. The temperature was reduced to 18 °C, and bacterial cultures were incubated overnight with agitation. Cells were harvested by centrifugation at 6000 × g for 20 minutes, and pellets were stored at −80°C for later use.

The pellet from 4 L of bacterial culture was resuspended in 60 mL of solution A (50 mM Tris–HCl pH 8.0, 500 mM NaCl, 20 mM imidazole) with 5 μL of benzonase endonuclease (Millipore, Sigma) and a tablet of protease inhibitor cocktail (cOmplete, EDTA-free, Roche). Continual cycles of 15 s ON and 59 s OFF at 60 % amplitude were performed for 5 min to lyse cells by sonication. The bacterial lysate was ultra-centrifuged at 125,748 × g for 1 hour. The soluble fraction was first passed through a 0.2 μm filter and then loaded using a peristaltic pump into a 5 ml HisTrap HP column equilibrated with solution A.

The column was connected to a fast protein liquid chromatography system (ÄKTA FPLC; GE Healthcare) and washed with solution A at 1 mL/min. When the absorbance at 280 nm stabilised near the baseline, the Tap3–Tke5 complex was eluted with 100% solution B (50 mM Tris–HCl pH 8, 500 mM NaCl and 500 mM imidazole) at 4 mL/min. Fractions of the central peak were pulled, and dithiothreitol (DTT) was added at a final concentration of 2 mM. The sample was then injected into a HiLoad Superdex 200 26/600 pg column, previously equilibrated with solution C (20 mM Tris–HCl pH 8, 150 mM NaCl and 2 mM DTT). The Tap3–Tke5 complex eluted as a single, monodispersed peak, and SDS-PAGE gel analysis showed two bands (**Fig. 1b**), which were confirmed by mass spectrometry to correspond to Tke5 and Tap3 (**Supplementary Data 1**). Central peak fractions corresponding to the complex were collected and concentrated using an Amicon centrifugal filter unit with a 10 kDa molecular mass cut-off (Millipore) to a final concentration of 9 mg mL^−1^ (approximately 1.5 mg of protein is obtained for each litter of culture).

About 0.5 ml of the freshly concentrated sample was injected into a Superdex™ 200 Increase 10/300 GL, previously equilibrated with solution C. Fractions of 50 µl were collected. The elution peak at 2.2 mg/ml was used to prepare grids after adding 0.05% CHAPS to the sample 5 minutes before vitrification, which is essential to avoid the preferred orientation pathology (Kampjut *et al*, 2021). Quantifoil^TM^ R 1.2/1.3 300 mesh grids were glow-discharged at a 0.41 mbar vacuum for 90 s. Then, 4 μL of purified Tap3– Tke5 at 2.2 mg/ml in solution C with 0.05% CHAPS were applied to the grids and blotted for 2.3 s in a Leica EM GP2 single-side blotting automated plunge freezer at 95% humidity. Grids were plunge-frozen in liquid ethane and stored in liquid nitrogen until data collection.

### Cryo-EM data collection and processing

#### Cryo-EM grid preparation

Grids were clipped in liquid nitrogen and were introduced into an in-house 300 kV Thermo-Fisher Titan Krios G4 transmission electron microscope, where data was collected paired with Gatan’s BioContinuum Imaging Filter. 19,112 movies were recorded on a K3 direct electron detector device at a nominal magnification of 130,000 x with a calibrated pixel size of 0.6462 Å. A defocus range of −0.8 to −2.5 μm was used, with a total dose of 48.8 e^−^/Å^2^ fractionated over 50 frames with a total exposure time of 1.3 s. Acquired image stacks were processed using CryoSPARC (Punjani *et al*, 2017) software, resulting in a final map that allowed the building of the atomic structure of Tap3–Tke5 at 2.8 Å.

#### Cryo-EM data processing

19,112 raw cryo-EM movies were preprocessed in CryoSPARC (Punjani *et al*, 2017) with Patch Motion Correction and Patch CTF Estimation. Template-based particle picking was performed with templates created from previous low-resolution reconstructions, followed by an initial 2D classification of 3× binned particles. Further rounds of 2D classification with varying parameters were used to improve particle quality and remove duplicates. The best 2D classes were re-extracted with no binning for an initial ab initio reconstruction into three classes. The best volume (referred to as the *initial volume*) was chosen for further 2D classification. From the best 2D classes, focused particle picking was then performed using Topaz (Bepler *et al*, 2020), with two separate models for different particle orientations, separating top/bottom views from side views. Particles were re-extracted with 3× binning (384 px box size, 128 px binning). Relaxed 2D classification criteria were applied to retain usable particles while removing the main unusable particles. Duplicates closer than 150 Å were removed when combining both particle sets. The final cleaning steps included the exclusion of aggregates. The initial volume was used as an input for Non-Uniform (NU) refinement (Punjani *et al*, 2020) with the latest particle set. A new ab initio reconstruction and heterogeneous refinement cycles were performed, in which the NU refinement volume coming from the initial volume was included as an input with the rest of the volumes from the ab initio reconstruction, and the best-resolved class was chosen for particle extraction with no binning and a box size of 384 px. A last heterogeneous refinement step was performed before re-extracting the particles in the best volume with a bigger box (600 px). Post-processing included Local CTF Refinement and Reference-Based Motion Correction. A final NU refinement was performed, achieving a 2.8 Å resolution. CryoSPARC (Punjani *et al*, 2017) Orientation Diagnostics was used to calculate the Sampling Compensation Factor (SCF*) and the corrected Fourier Amplitude Ratio (cFAR). SCF* and cFAR values of 0.827 and 0.57 indicate no particle orientation bias or map anisotropy. The final 2.8 Å density map was enhanced with DeepEMhancer (Sanchez-Garcia *et al*, 2021) for model building and visualisation. Cryo-EM data processing workflow and Cryo-EM maps and data quality are shown in **Supplementary Fig. 1, 2**. Data collection and processing statistics are provided in **Supplementary Table 1**.

### Model building and structure refinement

Approximately 65% of the Tke5 structure and 89% of the Tap3 structure were reconstructed using automatic *ab initio* model building with ModelAngelo (Jamali *et al*, 2024) and the density-modified DeepEMhancer (Sanchez-Garcia *et al*, 2021) cryo-EM map. The remaining regions were completed through multiple iterative rounds of manual model building performed in Coot (v0.9.8.91) (Casañal *et al*, 2020) and real-space-refinement in Phenix (1.20.1-4487 package) (Liebschner *et al*, 2019; Afonine *et al*, 2018) against the final NU refinement cryo-EM map. The final structure contains 94% and 92% of Tap3 and Tke5 sequences, respectively. Secondary structure and Ramachandran restraints were included in the refinement. MolProbity (Williams *et al*, 2018) and the PDB validation server (Gore *et al*, 2017) were used for model validation. A summary of the final model statistics is provided in **Supplementary Table 1**.

### Growth inhibition and immunity-driven recovery assays

*P. putida* KT2440 cells were electroporated with two compatible plasmids pS238D•M and pSEVA624C or their derivates harbouring *pelB·tke5*, *pelB·tke5_β*, *pelB·tke5_MIX, pelB·tke5_*α+β or *pelB·tke5_*α, and *tki5*, respectively, as described in **Supplementary Data 1**. Transformed cells were selected on LB-agar plates supplemented with Km 50 µg mL^-1^ (for pS238D•M) and Gm 25 µg mL^-^ (for pSEVA624C). The presence of the plasmid was confirmed by colony PCR using primers P10-P11 for pS238D•M derivatives and P12-P13 for pS624C derivatives, as listed in **Supplementary Data 1**.

For toxicity assays, overnight cultures of each strain were grown in an LB medium supplemented with the corresponding antibiotics. The following day, cultures were diluted to an OD600 of 0.05 in fresh medium, and 1 mM of IPTG was added to induce *tki5*. Aliquots of 200 μL were added to a microtiter plate, and OD600 values were measured every 15 min at 30 °C using the Synergy/H1 microplate reader (Biotek). After an initial two-hour incubation, 0.5 mM of 3-methylbenzoate (3-*m*Bz) was added to each well and cells were incubated for another 12 h. Data was collected for three biological replicates, each with technical duplicates.

### Bioinformatic and computational analysis

#### Tke5 topology prediction

The topology of Tke5 was predicted using the TMHMM-2.0 algorithm (DTU Health Tech (Krogh *et al*, 2001)) with the default settings.

#### Structural-base homology search of Tap3 and Tke5

The cryo-EM structures of Tke5 and Tap3 were used as a query to search for homologues using Foldseek Search Tool (van Kempen *et al*, 2024).

#### Tke5 protein structure prediction and membrane positioning

The C-terminal residues (608-996) of Tke5 (Tke5^608-996^) were modelled using AlphaFold 3 (Abramson *et al*, 2024). The disordered regions in the cryo-EM Tap3–Tke5 structure, including a loop within the putative receptor-binding domain (RBD) (residues 886-909), were modelled using AlphaFold_unmasked (Mirabello *et al*, 2024). The spatial orientation and optimal positioning of the Tke5 pre-pore state and the AlphaFold 3 prediction of Tke5^608-996^ in a model Gram-negative inner membrane were predicted using the PPM 3.0 server (Protein Property Prediction and Modelling Server (Lomize *et al*, 2022)). The model membrane composition is PE:PG:CL (79:19:2) containing acyl chains 15:0, 16:0, 16:1, 18:1, cy17:0. PPM 3.0 determines the energetically optimal spatial position of a protein structure by minimising its free energy of transfer from water to the membrane environment, treating the membrane as a fluid anisotropic solvent.

### Statistical analyses

Statistical analysis of the growth inhibition and immunity-driven recovery assays was performed using GraphPad Prism 9 version 9.5.1. Details regarding specific statistical tests and data interpretation are provided in the corresponding figure legend.

## Data availability

The authors declare that source data supporting the findings of this study are available within the paper and its supplementary information files. The Supplementary Information contains **Supplementary Fig. 1-11** and **Supplementary Table 1-4.** Atomic coordinates of the Tap3–Tke5 structure have been deposited in the Protein Data Bank (PDB) (accession PDB code id: 9R8G). The cryo-EM map is available from the Electron Microscopy Data Bank (accession code id: EMD-53820). **Supplementary Data 1** is available as a supplementary file and contains a list of strains, plasmids, primers, materials and the LC-ESI-MS report of the purified Tap3–Tke5 complex. Uncropped and unedited gel images are included in **Supplementary Fig. 11**.

## Structure visualisation tools

Molecular graphics and analyses were performed using UCSF Chimera (Pettersen *et al*, 2004) and UCSF ChimeraX (Goddard *et al*, 2018).

## Acknowledgements

We acknowledge the Basque Resource for Electron Microscopy (BREM) for access to their cryo-EM facility and their staff for data-acquisition support. The authors would like to thank Diamond Light Source for sample screening and data collection access and support of the cryo-EM facilities at the UK’s national Electron Bio-imaging Centre (eBIC) [under proposals EM BI38262], funded by the Wellcome Trust, MRC and BBSRC. This study was supported by the Spanish Ministry of Science and Innovation (MCIN/AEI/10.13039/501100011033/ FEDER, UE projects PID2021-123000OB-I00, CNS2022-135585 to PB, and project PID2021-127816NB-I00 to DAJ) and the Basque Government (IT1745-22 to DAJ). The funders had no role in study design, data collection and analysis, decision to publish, or preparation of the manuscript.

---

**Movie 1 Movie showing the 2.8 Å cryo-EM map of Tap3-Tke5 spinning along the *y*-axis of reference.**

