## Supplementary Information for "The cryo-EM structure of an adaptor-effector complex reveals the mechanism of a widespread pore-forming toxin family"

**This file includes:**

Supplementary Fig. 1-11

Supplementary Table 1-4

Supplementary Data 1 information

### SUPPLEMENTARY INFORMATION FIGURES

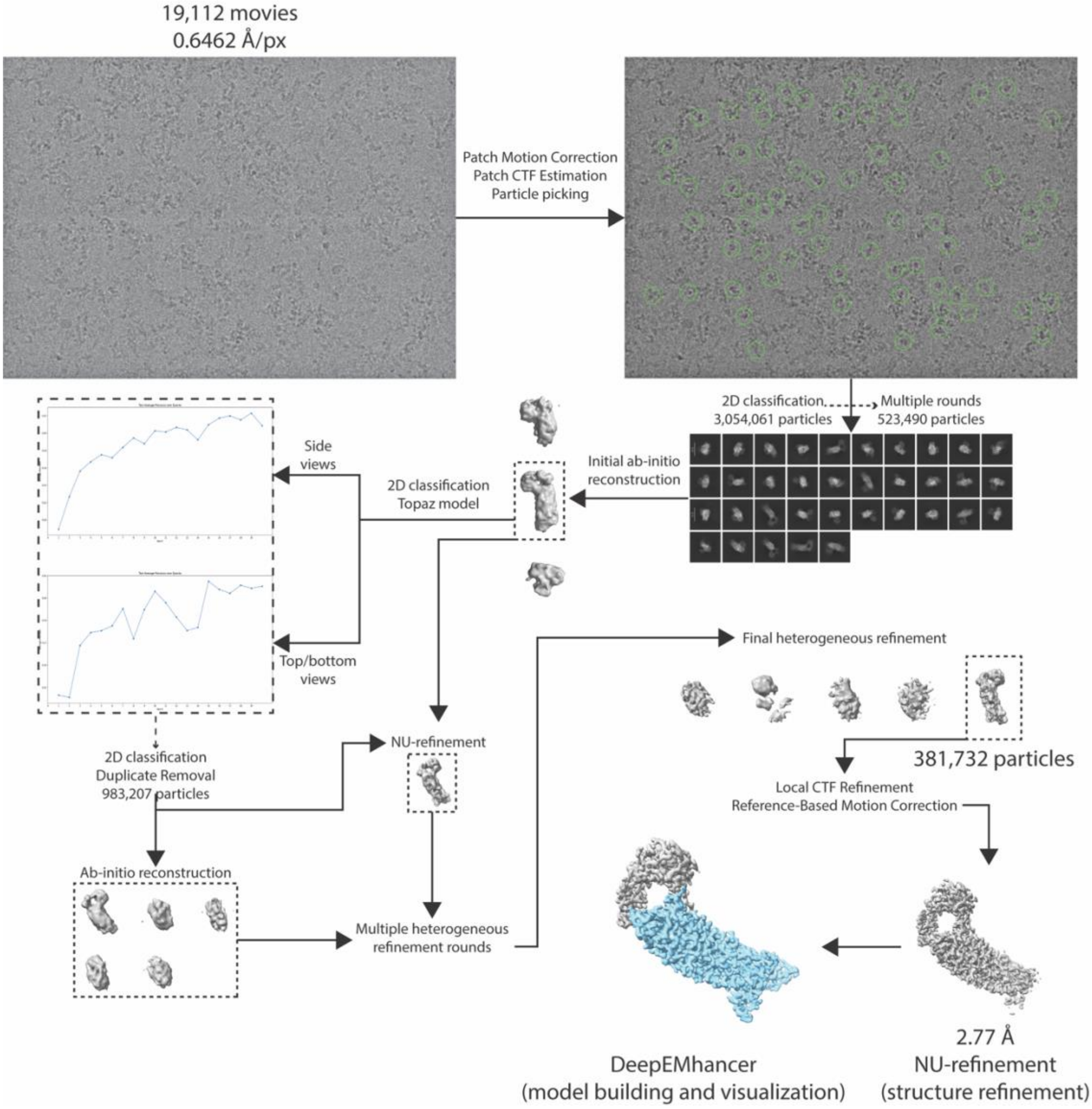


**Supplementary Figure 1. Cryo-EM data processing workflow for the *Pseudomonas putida* Tke5 toxin.** Workflow encompassing data acquisition through to the reconstruction of the final 2.8 Å resolution map. Cryo-EM movies were processed in CryoSPARC v.4.5.3[1]. Initial particle picking was refined through 2D classification iterations, followed by ab initio reconstruction and selection of the best volume. Focused picking with Topaz[2], along with further 2D classifications, improved both particle quantity and quality. Five new ab initio volumes were generated using the new particles, and the best volume from the previous ab initio was refined with the updated particle set. All six volumes were included in subsequent heterogeneous refinement cycles. The best class from the final round of heterogeneous refinement was selected for post-processing. The fully solved density map was further enhanced using DeepEMhancer[3].

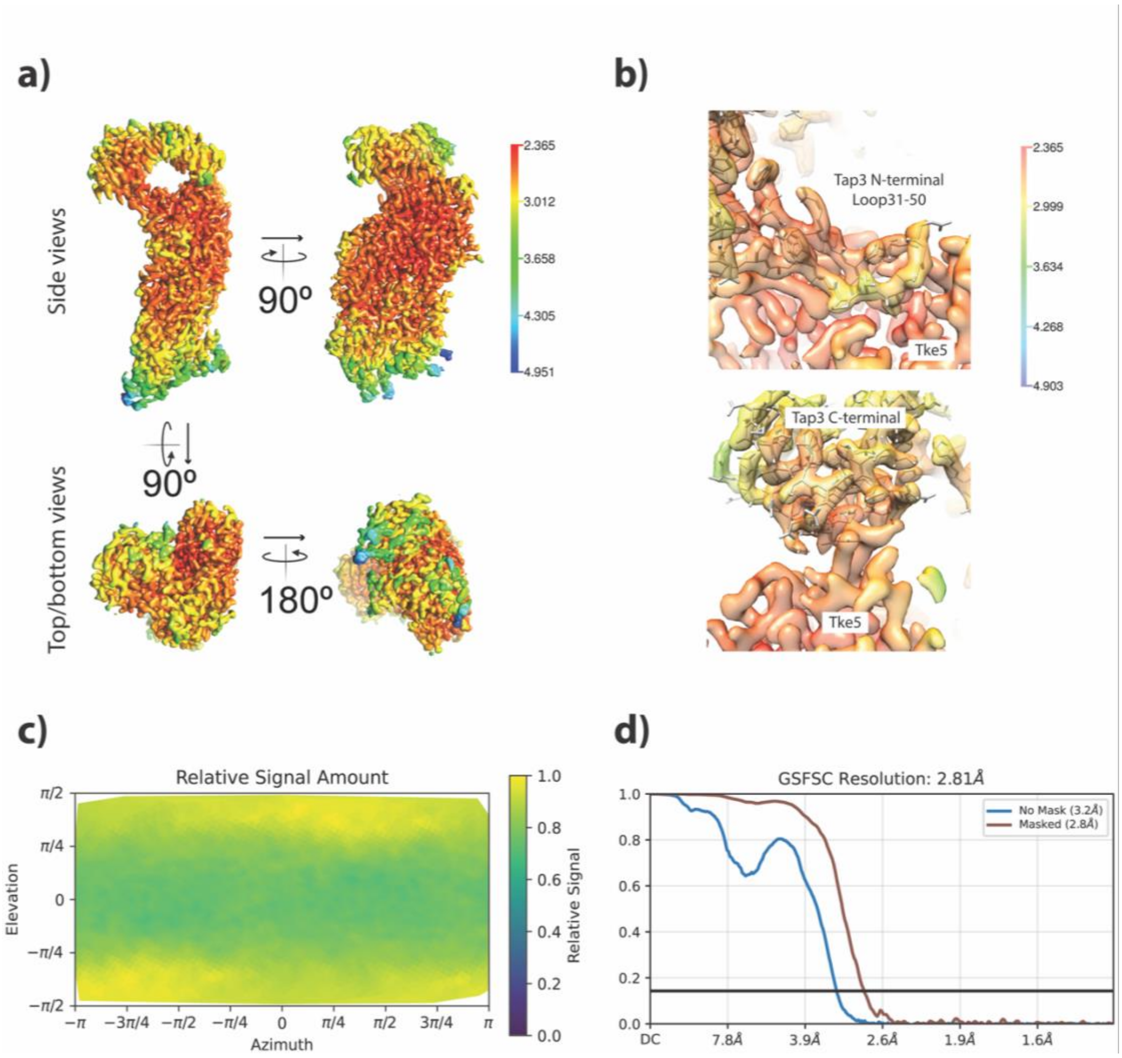


**Supplementary Figure 2. Cryo-EM maps and data quality a)** Local resolution of the Tap3-Tke5 EM map, represented in the surface as a gradient with UCSF Chimera[4]. Different side and top/bottom views are shown. **b)** Zoomed local resolution EM map in the N-terminal Loop$^{31\text{-}50}$ and the C-terminal of Tap3 residues (atomic stick representation) interacting with Tke5 residues (no structure is shown, only the cryo-EM map). **c)** Relative signal amount plot of the EM map. CryoSPARC Orientation Diagnostics was used to calculate the Sampling Compensation Factor (SCF*) and the corrected Fourier Amplitude Ratio (cFAR). SCF* and cFAR values of 0.827 and 0.57 indicate no orientation bias or anisotropy. **d)** Global resolution estimation of the EM map. Gold-standard Fourier shell correlation (GSFSC) curves were calculated with and without masking. The masked resolution is estimated at 2.8 Å using a 0.143 cutoff criterion.

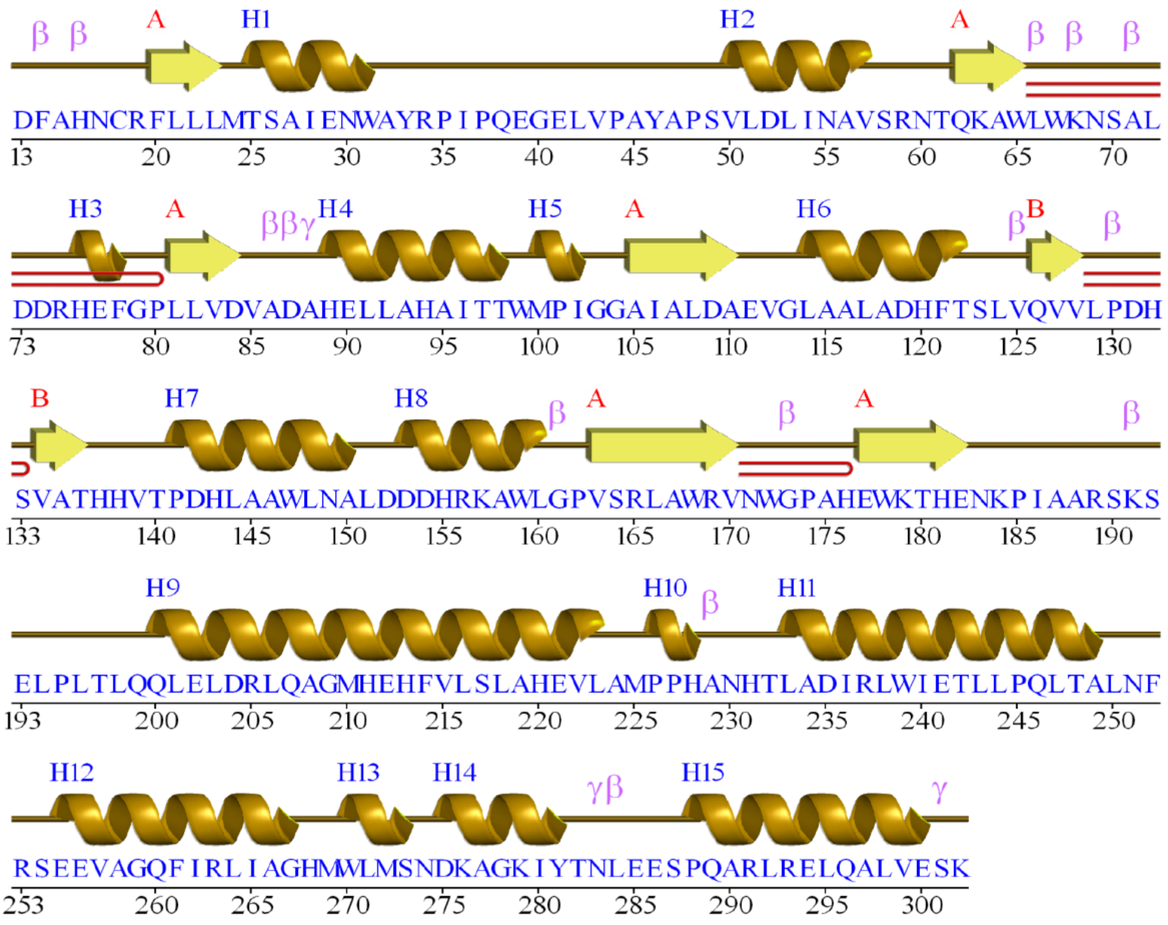


**Supplementary Fig. 3.** Secondary structure diagram of Tap3 (290 modelled residues). The diagram was generated using PDBsum[5]. It displays the secondary structure elements, including 15 α-helices, in gold, and 2 β-sheets (A, with 7 β-strands; and B, with 2 β-strands), shown in yellow. Additionally, the diagram highlights β-turns (14), γ-turns (3), and β- hairpins (3).

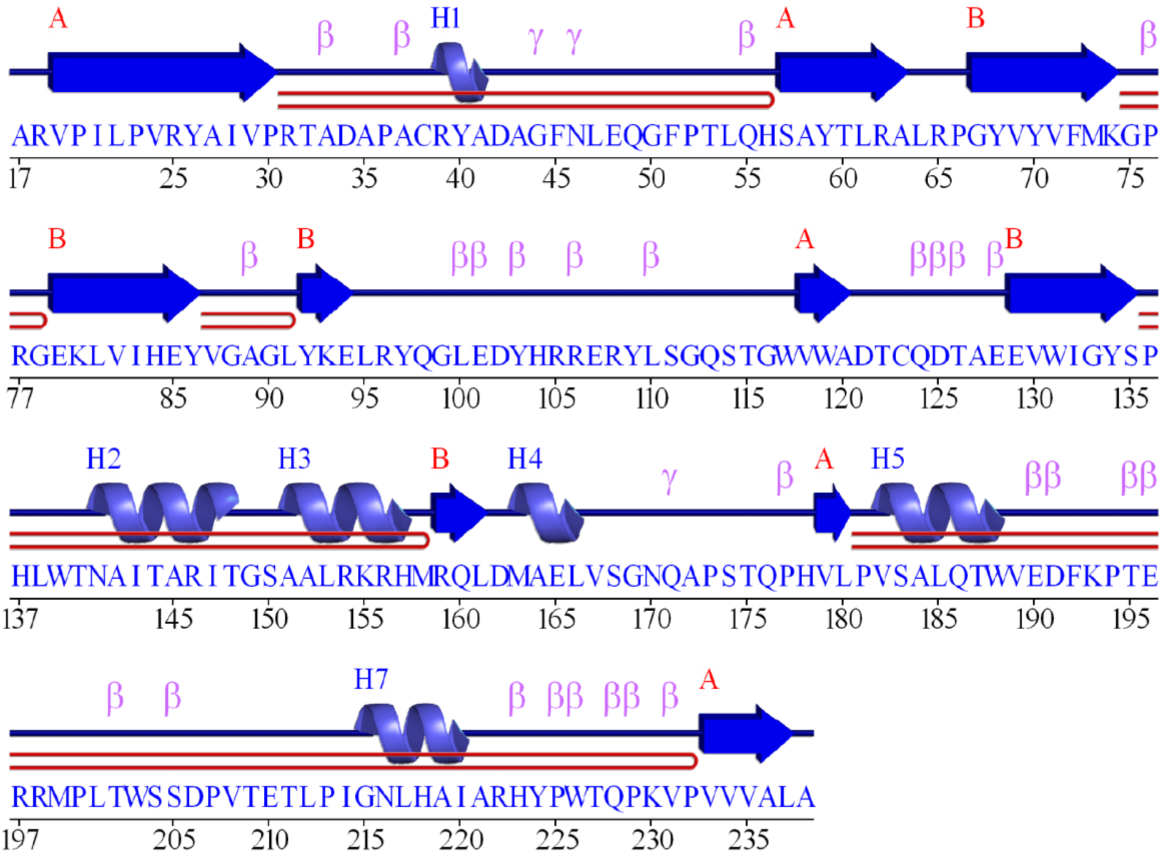


**Supplementary Fig. 4.** Secondary structure diagram of MIX (222 modelled residues). The diagram was generated using PDBsum. It displays the secondary structure elements, including 7 α-helices, in light blue, and 2 β-sheets (A, with 5 β-strands; and B, with 5 β-strands), shown in dark blue. Additionally, the diagram highlights β-turns (27), γ-turns (3), and β-hairpins (5).

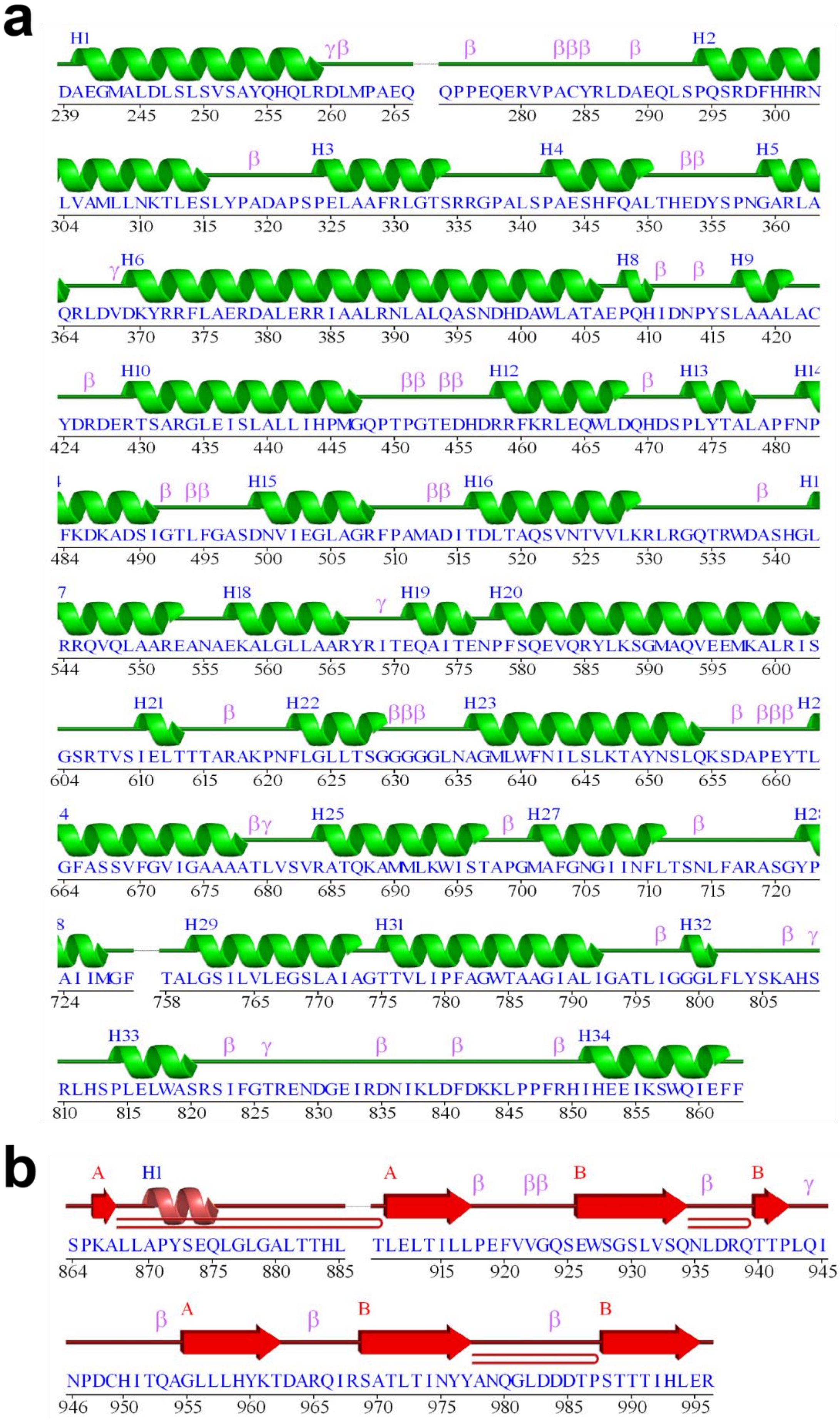
a
H1 γβ β βββ β H2
DAEGMALDLSLSVSAYQHQLRDLMPAEQ QPPEQERVPACYRLDAEQLSPQSRDFHHRN
239 245 250 255 260 265 280 285 290 295 300
β H3 H4 ββ H5
LVAMLLNKTLESLYPADAPSPELAAFRLGTSRRGPALSPAESHFQALTHEDYSPNGARLA
304 310 315 320 325 330 335 340 345 350 355 360
γ H6 H8 β β H9
QRLDVDKYRRFLAERDALERRIAALRNLALQASNDHDAWLATAEPQHIDNPYSLAAALAC
364 370 375 380 385 390 395 400 405 410 415 420
β H10 ββ ββ H12 β H13
YDRDERTSARGLEISLALLIHPMGQPTPGTEDHDRRFKRLEQWLDQHDSPLYTALAPFNP
424 430 435 440 445 450 455 460 465 470 475 480
β ββ H15 ββ H16 β
FKDKADSIGTLFGASDNVIEGLAGRFPAMADITDLTAQSVNTVVLKRLRGQTRWDASHGL
484 490 495 500 505 510 515 520 525 530 535 540
H18 γ H19 H20
RRQVQLAAREANAEKALGLLAARYRITEQAITENPFSQEVQRYLKSGMAQVEEMKALRIS
544 550 555 560 565 570 575 580 585 590 595 600
H21 β H22 βββ H23 β βββ
GSRTVSIELTTTARAKPNFLGLLTSGGGGGLNAGMLWFNILSLKTAYNSLQKSDAPEYTL
604 610 615 620 625 630 635 640 645 650 655 660
βγ H25 β H27 β
GFASSVFGVIGAAAATLVSVRATQKAMMLKWISTAPGMAFGNGIINFLTSNLFARASGYP
664 670 675 680 685 690 695 700 705 710 715 720
H29 H31 β H32 β γ
AIIMGF TALGSILVLEGSLAIAGTTVLIPFAGWTAAGIALIGATLIGGGLFLYSKAHS
724 758 765 770 775 780 785 790 795 800 805
H33 β γ β β β H34
RLHSPLELWASRSIFGTRENDGEIRDNIKLDFDKKLPPFRHIHEEIKSWQIEFF
810 815 820 825 830 835 840 845 850 855 860
b
A H1 A β ββ B β B γ
SPKALLAPYSEQLGLGALTTHL TLELTILLPEFVVGQSEWSGSLVSQNLDRQTTPLQI
864 870 875 880 885 915 920 925 930 935 940 945
β A β B β B
NPDCHITQAGLLLHYKTDARQIRSATLTINYYANQGLDDDTPSTTTIHLER
946 950 955 960 965 970 975 980 985 990 995

**Supplementary Fig. 5.** Secondary structure diagram of the α+ β-domain. **(a)** The α-region is shown (590 modelled residues), and **(b)** the β-rich region is shown (109 modelled residues). The diagram was generated using PDBsum. **(a)** It displays the secondary structure elements, including 34 α-helices, shown in green. Additionally, the diagram highlights β-turns (40) and γ-turns (7). See Supplementary Table 4 for more details, including predicted TM helices. **(b)** It displays the secondary structure elements, including 1 α-helices, in light red, and 2 β-sheets (A, with 3 β-strands; and B, with 4 β-strands), shown in dark red. Additionally, the diagram highlights β-turns (7), γ-turns (1), and β-hairpins (3).

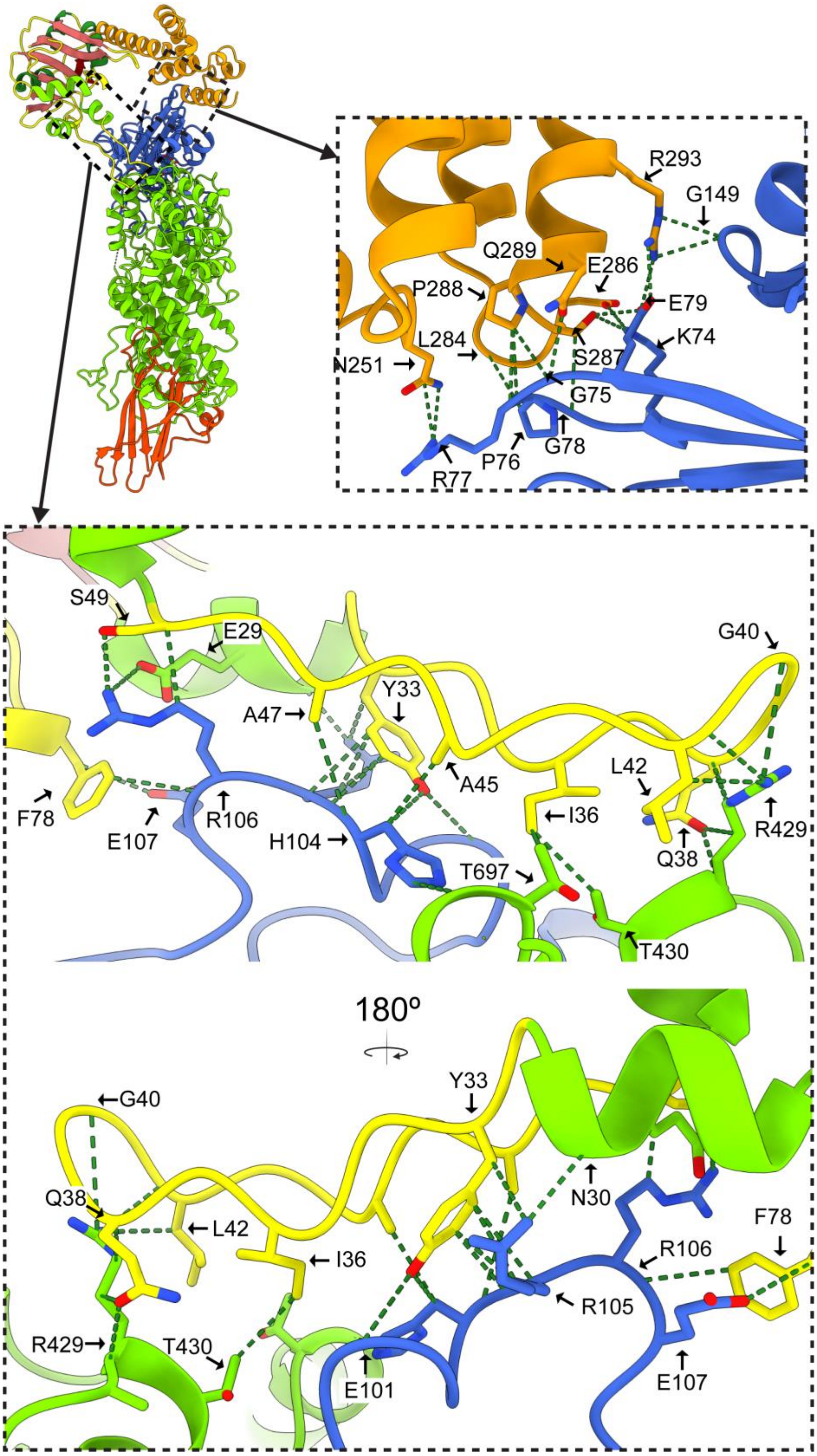


**Supplementary Figure 6. All hydrogen bonds between Tap3 and Tke5.** An overview of the structure is shown using the same colour scheme as in Fig. 3, with quadrants indicated for zoomed-in views. The upper quadrant highlights the interaction between the α-helical bundle of Tap3 (in orange) and the MIX domain (in blue). The lower quadrant focuses on the interaction between the Tap3-Loop$^{31-50}$ (in yellow) and both the MIX domain and the α-domain (in green), shown with a 180° rotation to reveal all hydrogen bond interactions. Interactions were visualised in ChimeraX[6]. For a detailed list of these interactions, see Supplementary Table 2.

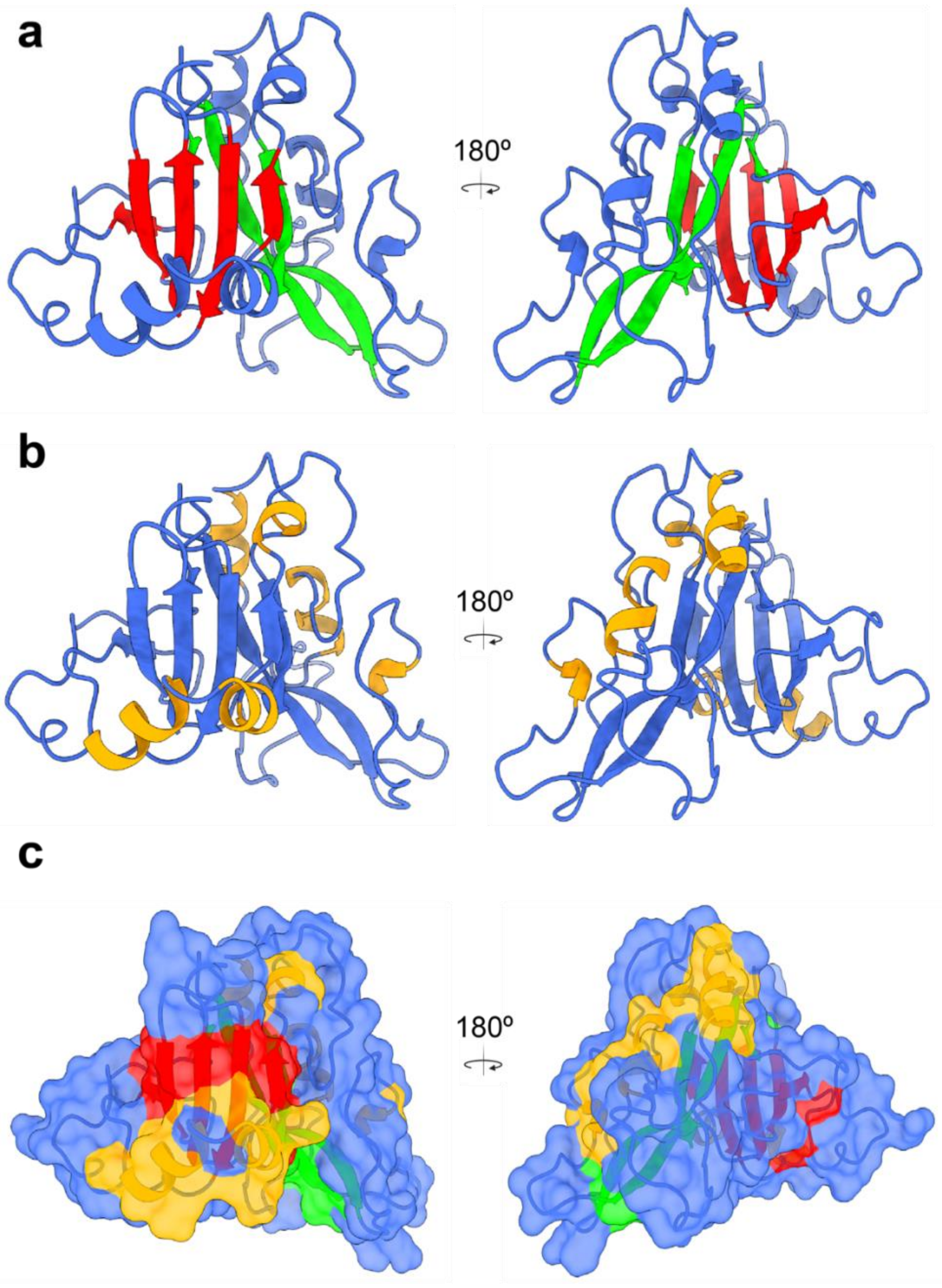


**Supplementary Figure 7. The structural fold of the Tke5 MIX domain.** Two views of the MIX domain (residues 1-238) are shown, rotated 180º for optimal visualisation. **(a).** The two central β-sheets are distinctly highlighted, with one β-sheet coloured in red and the other coloured in green, both composed of five β-strands. **(b).** The six α-helices are displayed in orange, showing their spatial arrangement relative to the central β-sheets. **(c).** The secondary structural elements maintain the same colour scheme as in panels (a) and (b), with the addition of the MIX domain's molecular surface depicted in blue. This surface representation clearly shows the characteristic pyramid-like architecture formed by the interplay of the β-sheets and α-helices within the MIX fold.

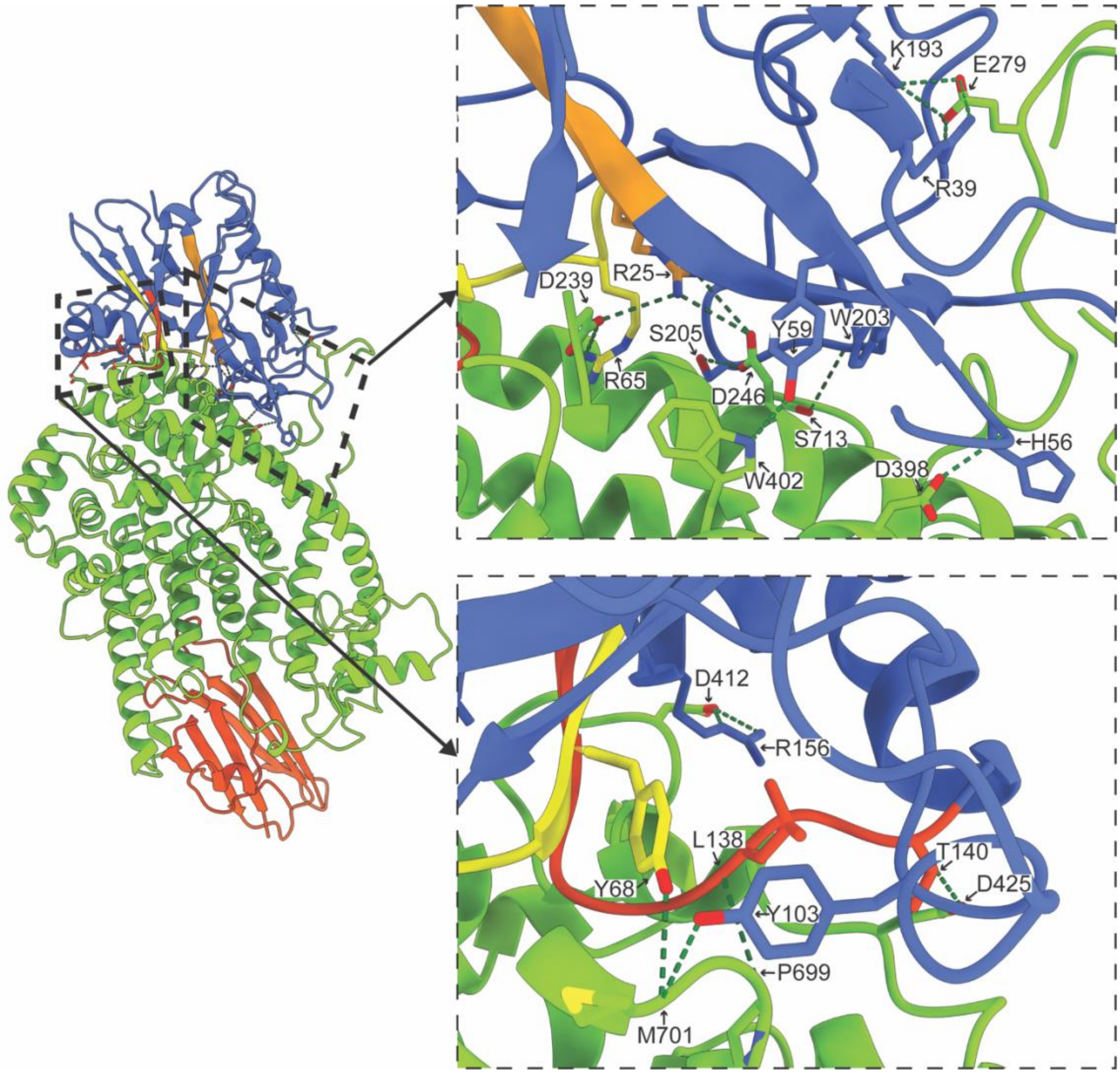


**Supplementary Figure 8.** Hydrogen bonds between the MIX domain (shown in blue; MIX motif highlighted in red, yellow, and orange) and the α+β domain (shown in green) are displayed. For a detailed list of these interactions, see Supplementary Table 2.

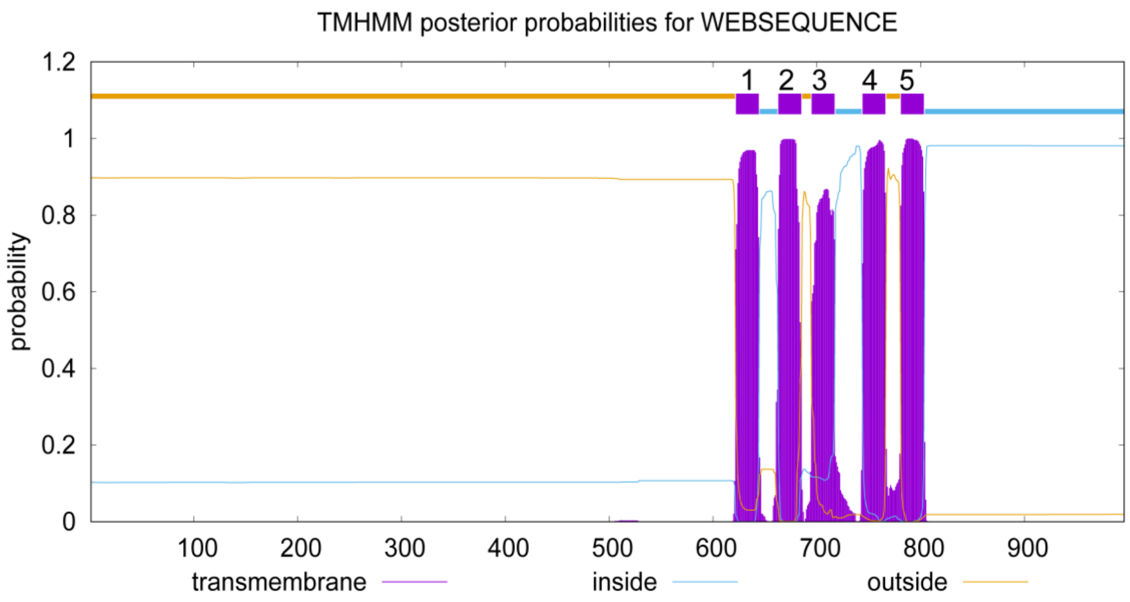


TMH1: 622-644 TMH2: 663-685 TMH3: 695-717 TMH4: 744-766 TMH5: 781-803

**Supplementary Figure 9. Transmembrane helices predicted for Tke5 using TMHMM v2.0 based on a Hidden Markov Model.** This plot displays posterior probabilities for each amino acid residue being located in a transmembrane helix (TM helix), on the inside (cytoplasmic side), or the outside (non-cytoplasmic side) of the membrane. The predicted topology generated via the TMHMM server[7] is as follows: outside (residues 1-621), TMH1 (622-644), inside (645-662), TMH2 (663-685), outside (686-694), TMH3 (695-717), inside (718-743), TMH4 (744-766), outside (767-780), TMH5 (781-803), and inside (804-996).

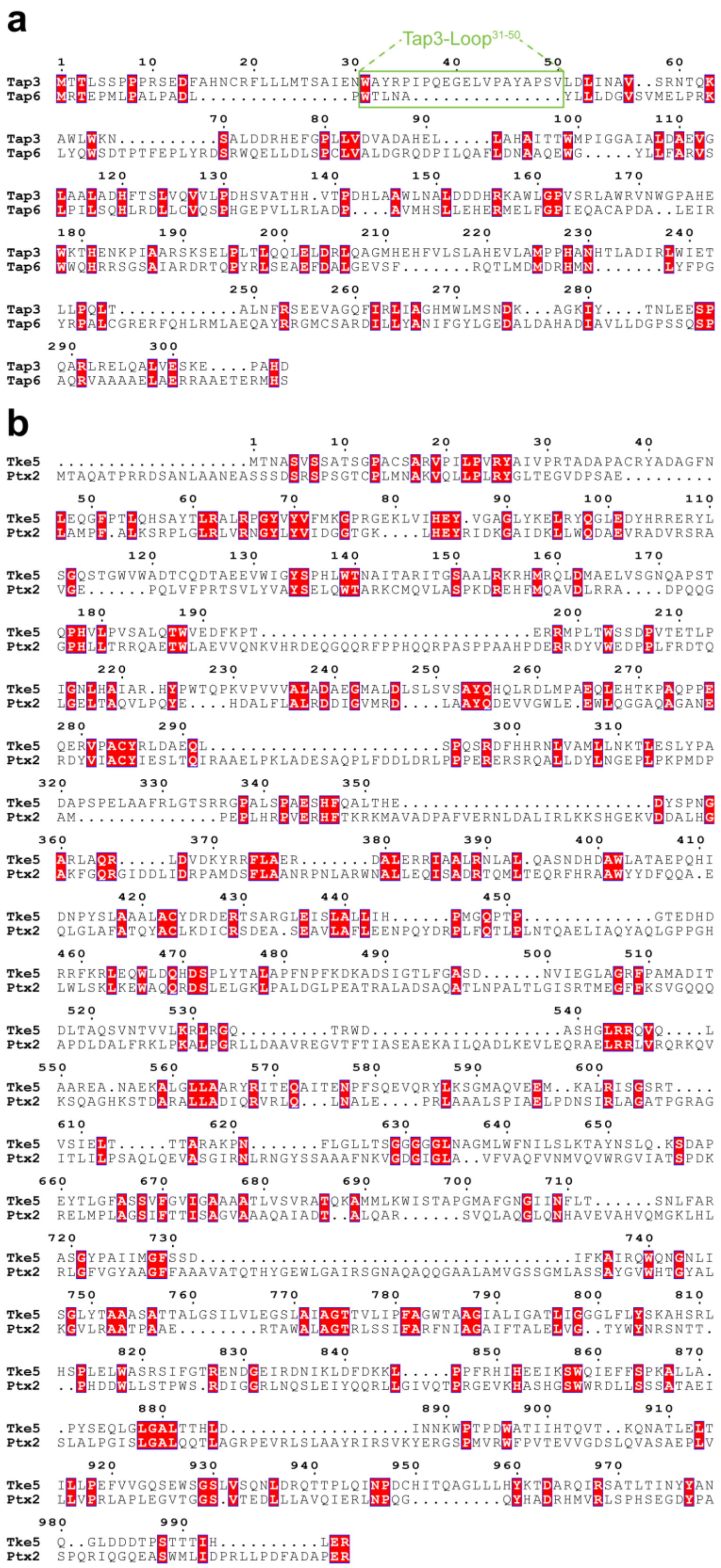


**Supplementary Figure 10. Sequence alignment of a) *Pp*Tap3 and *Pa*Tap6 and b) *Pp*Tke5 and *Pa*Ptx2.** The alignment was generated using Clustal Omega[8] and visualised with ESPript 3.0[9]. The Strict similarity colouring scheme was applied to highlight strictly conserved residues. The sequence identity shared by the aligned sequences is 16 % for a) and 17 % for b). In the case of a), the Tap3-Loop$^{31\text{-}50}$ insertion is boxed in green, while Tap6 shows a gap in that region.

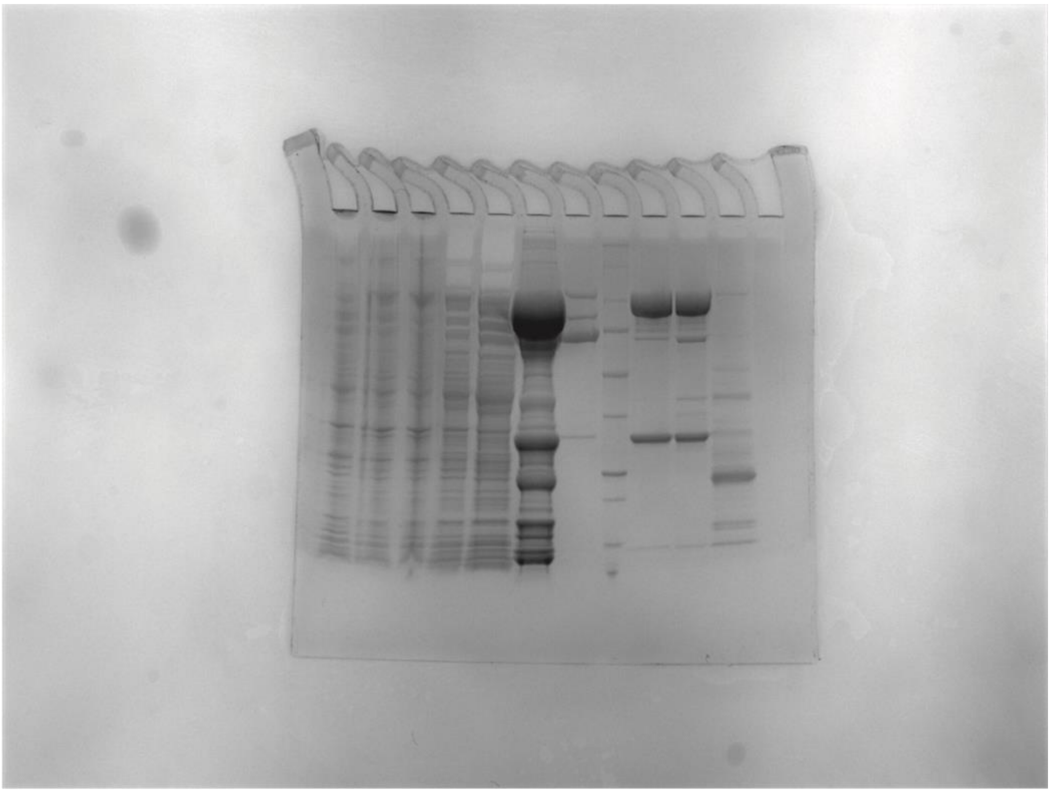

**Supplementary Figure 11. Uncropped and unedited SDS-gel image.** Representative sodium dodecyl sulphate-polyacrylamide gel electrophoresis (SDS-PAGE) of Tap3-Tke5 heterologous expression and its purification. Lane 1, non-induced fraction; Lane 2, induced fraction; Lane 3, insoluble fraction; Lane 4, soluble fraction; lane 5, flowthrough from 5 ml His-Trap column; lane 6, elution from 5 ml His-Trap column; lane 7, first small peak in the HiLoad Superdex 200 26/600 pg; lane 8, molecular weight standard (Precision Plus Protein$^{TM}$ Standards, BIO-RAD); lanes 9 and 10, central peak in the HiLoad Superdex 200 26/600 pg that corresponds to Tap3-Tke5 complex; lane 11, third small peak in the HiLoad Superdex 200 26/600 pg. The identity of the Tke5 (around 110 kDa) and Tap3 band (around 35 kDa) was confirmed by mass spectrometry (Supplementary Data 1).

#### SUPPLEMENTARY TABLES

**Supplementary Table 1.** Cryo-EM data collection, data processing and model statistics.

| **Data Collection** | **EMD 53820, PDB 9R8G** |
|---|---|
| Microscope | ThermoFisher Titan Krios |
| Magnification | 130,000 × |
| Voltage (kV) | 300 |
| Camera | Gatan K3 |
| Pixel size (Å/pixel) | 0.6462 |
| Total electron dose ($e^{-}$/Å$^2$) | 48.8 |
| Exposure time (s) | 1.3 |
| Defocus range ($\mu$m) | −0.8 to −2.5 |
| Number of images | 19112 |
| **Processing** | |
| Number of select Micrographs | 18617 |
| Number of initial particles | 3054061 |
| Number of final particles | 381732 |
| Final resolution (Å) | 2.8 |
| Symmetry | C1 |
| Map sharpening B factor (Å$^2$) | 100.8 |
| **Orientation Diagnostics** | |
| SCF* | 0.827 |
| cFAR | 0.57 |
| **Structure Composition /Validation** | |
| No. of chains | 2 |
| Non-hydrogen atoms | 8811 |
| Protein residues | 1211 |
| Bond RMSD lengths (Å) (#>4 $\sigma$) | 0.004 (0) |
| Bond RMSD angles (°) (#>4 $\sigma$) | 0.452 (3) |
| Molprobity score | 1.15 |
| Clashcore | 3.42 |
| Rotamers outliers (%) | 0.12 |
| Ramachandran favored (%) | 97.92 |
| Ramachandran allowed (%) | 2.08 |
| Ramachandran outliers (%) | 0 |
| CC (mask) | 0.86 |
| CC (box) | 0.87 |
| CC (peaks) | 0.78 |
| CC (volume) | 0.85 |

**Supplementary Table 2.** Salt-bridge and hydrogen bond interactions between Tap3 and Tke5. Interactions were computed in PDBsum.

| **Tke5** | **Tap3** | **Distance (Å)** |
|---|---|---|
| Salt-bridge interactions | | |
| LYS74 | GLU286 | 2.93 |
| GLU79 | ARG293 | 3.04 |
| ARG106 | GLU29 | 2.91 |
| ARG106 | GLU77 | 3.45 |
| Hydrogen bond interactions | | |
| LYS74 | GLU286 | 2.93 |
| LYS74 | SER287 | 2.99 |
| GLU79 | ARG293 | 3.06 |
| GLU79 | SER287 | 2.76 |
| GLU79 | ARG293 | 3.04 |
| GLU101 | TYR33 | 2.76 |
| ARG105 | ASN30 | 3.13 |
| ARG106 | GLU29 | 2.91 |
| ARG429 | GLN38 | 2.94 |
| ARG429 | GLY40 | 3.07 |

**Supplementary Table 3.** Salt-bridge and hydrogen bond interactions between Tke5 MIX domain and α+β domain. Interactions were computed in PDBsum.

| MIX domain | α+β domain | Distance (Å) |
|---|---|---|
| Salt-bridge interactions | | |
| ARG25 | ASP239 | 3.80 |
| ARG39 | GLU279 | 3.39 |
| ARG65 | ASP239 | 2.83 |
| ARG156 | ASP412 | 3.30 |
| ARG193 | GLU279 | 3.06 |
| Hydrogen bond interactions | | |
| HIS56 | ASP398 | 2.86 |
| TYR59 | TRP402 | 2.80 |
| ARG65 | ASP239 | 2.83 |
| TYR68 | MET701 | 3.10 |
| TYR103 | MET701 | 3.10 |
| LEU138 | PRO699 | 3.02 |
| THR140 | ASP425 | 2.78 |
| ARG156 | ASP412 | 3.30 |
| LYS193 | GLU279 | 3.06 |
| TRP203 | SER713 | 2.74 |
| SER205 | ASP246 | 3.35 |
| SER205 | ASP246 | 2.75 |

**Supplementary Table 4.** Table showing the 34 helices α-helices of the α-region of Tke5. Nº (number), S (start), E (end), T (Type), R (number of residue), L (length), Ur (unit rise), RpT (residues per turn), P (pitch), D (deviation from ideal), S (sequence). Predicted TM Helices on the surface are colored in blue, and TM Helices on the core are colored in yellow. This table was built with PDBsum.

| Nº | S | E | T | R | L | Ur | RpT | P | D | S |
|---|---|---|---|---|---|---|---|---|---|---|
| 1. | A240 | R259 | H | 20 | 29.73 | 1.47 | 3.57 | 5.25 | 10.2 | AEGMALDLSLSVSAYQHQL R |
| 2. | P294 | S315 | H | 22 | 33.10 | 1.48 | 3.67 | 5.44 | 14.2 | PQSRDFHHRNLVAMLLNKT LES |
| 3. | P324 | S334 | H | 11 | 16.73 | 1.48 | 3.67 | 5.43 | 6.0 | PELAAFRLGTS |
| 4. | P342 | L350 | H | 9 | 13.76 | 1.48 | 3.57 | 5.29 | 7.4 | PAESHFQAL |
| 5. | G359 | Q364 | H | 6 | 10.11 | 1.59 | 3.51 | 5.60 | 8.7 | GARLAQ |
| 6. | D369 | L403 | H | 35 | 49.45 | 1.43 | 6.41 | 9.19 | 66.2 | DKYRRFLAERDALERRIAAL RNLALQASNDHDAWL |
| 7. | A404 | A406 | G | 3 | - | - | - | - | - | ATA |
| 8. | P408 | H410 | G | 3 | - | - | - | - | - | PQH |
| 9. | L417 | L421 | H | 5 | 8.07 | 1.51 | 3.58 | 5.39 | 21.2 | LAAAL |
| 10. | R429 | I443 | H | 15 | 22.19 | 1.46 | 3.68 | 5.36 | 7.8 | RTSARGLEISLALLI |
| 11. | H444 | G447 | G | 4 | 7.39 | 1.86 | 3.40 | 6.32 | 41.2 | HPMG |
| 12. | R458 | D468 | H | 11 | 17.42 | 1.55 | 3.57 | 5.53 | 8.0 | RRFKRLEQWLD |
| 13. | P473 | L478 | H | 6 | 9.58 | 1.49 | 3.46 | 5.14 | 4.9 | PLYTAL |
| 14. | N482 | I491 | H | 10 | 15.33 | 1.48 | 3.63 | 5.38 | 1.4 | NPFKDKADSI |
| 15. | D499 | R508 | H | 10 | 15.70 | 1.50 | 3.63 | 5.45 | 3.7 | DNVIEGLAGR |
| 16. | T516 | K529 | H | 14 | 21.91 | 1.51 | 3.78 | 5.72 | 11.4 | TDLTAQSVNTVVLK |
| 17. | L543 | E553 | H | 11 | 17.11 | 1.51 | 3.60 | 5.44 | 7.9 | LRRQVQLAARE |
| 18. | E557 | R566 | H | 10 | 15.42 | 1.49 | 3.61 | 5.38 | 1.8 | EKALGLLAAR |
| 19. | E571 | E576 | H | 6 | 9.50 | 1.52 | 3.53 | 5.35 | 5.4 | EQAITE |
| 20. | P578 | S603 | H | 26 | 38.38 | 1.46 | 3.64 | 5.32 | 11.9 | PFSQEVQRYLKSGMAQVE EMKALRIS |
| 21. | I610 | T613 | H | 4 | 6.84 | 1.66 | 3.49 | 5.81 | 40.7 | IELT |
| 22. | F622 | G629 | H | 8 | 10.81 | 1.39 | 3.90 | 5.43 | 13.6 | FLGLLTSG |
| 23. | A636 | Q654 | H | 19 | 28.51 | 1.47 | 3.63 | 5.34 | 7.5 | AGMLWFNILSLKTAYNSLQ |
| 24. | L663 | A678 | H | 16 | 24.56 | 1.50 | 3.61 | 5.41 | 7.3 | LGFASSVFGVIGAAAA |
| 25. | R684 | W694 | H | 11 | 17.03 | 1.53 | 3.57 | 5.47 | 11.8 | RATQKAMMLKW |

| | | | | | | | | | | |
|---|---|---|---|---|---|---|---|---|---|---|
| 26. | I695 | T697 | G | 3 | - | - | - | - | - | IST |
| 27. | M701 | L711 | H | 11 | 17.11 | 1.52 | 3.59 | 5.44 | 7.7 | MAFGNGIINFL |
| 28. | Y722 | M727 | H | 6 | 10.30 | 1.62 | 3.58 | 5.81 | 7.6 | YPAIIM |
| 29. | L760 | S769 | H | 10 | 15.89 | 1.54 | 3.55 | 5.48 | 5.9 | LGSILVLEGS |
| 30. | L770 | A773 | G | 4 | 6.80 | 1.49 | 3.93 | 5.86 | 41.7 | LAIA |
| 31. | T775 | I792 | H | 18 | 26.91 | 1.46 | 3.69 | 5.37 | 11.2 | TTVLIPFAGWTAAGIALI |
| 32. | G799 | L801 | G | 3 | - | - | - | - | - | GGL |
| 33. | P814 | S820 | H | 7 | 10.87 | 1.53 | 3.54 | 5.40 | 12.2 | PLELWAS |
| 34. | I851 | F862 | H | 12 | 19.44 | 1.57 | 3.54 | 5.56 | 10.3 | IHEEIKSWQIEF |

#### SUPPLEMENTARY DATA

**Supplementary Data 1. Strains, plasmids, oligonucleotides primers, materials, and Mass spectrometry of Tap3-Tke5**

**Sheet 1.** Bacterial strains were used in this study. Table showing each strain (A), with its relevant characteristics (B), use (C) and source (D). The antibiotic resistance marker is identified as follows: Rif, rifampicin. **Sheet 2.** Plasmids used in this study. Table showing each plasmid (A), with its relevant characteristics (B), use (C), source (D), DNA sequence (E), and target protein sequence (F). The antibiotic resistance markers are identified as follows: Amp, ampicillin; Km, kanamycin; Gm, gentamicin; Sm, streptomycin. **Sheet 3.** Oligonucleotide primers used in this study. Table showing its primer number (A), its description (B), such as restriction enzyme used for cloning and primer orientation (forward or F and reverse or R), and its (5' - 3') sequence (C). **Sheet 4.** Relevant materials used in this study. Table showing the name (A), company (B), reference (C) and application (D) of a list of chemicals and equipment. **Sheet 5.** Mass spectrometry identification of Tke5 and Tap3 pure bands. Table showing the proteins identified by mass spectrometry with their accession number (A), description (B), score (C), coverage (D), proteins (E), unique peptides (F), peptides (G), PSMs (H), number of amino acids (I), molecular weight (J) and isoelectric point (K).
